# Cell Atlas at Single-Nuclei Resolution of the Adult Human Adrenal Gland and Adrenocortical Adenomas

**DOI:** 10.1101/2022.08.27.505530

**Authors:** Barbara Altieri, A. Kerim Secener, Somesh Sai, Cornelius Fischer, Silviu Sbiera, Panagiota Arampatzi, Sabine Herterich, Laura-Sophie Landwehr, Sarah N. Vitcetz, Caroline Braeuning, Martin Fassnacht, Cristina L. Ronchi, Sascha Sauer

## Abstract

The human adrenal gland is a complex endocrine tissue. Developmental studies on this tissue have been limited to animal models or human foetus. Here, we present a cell atlas analysis of the adult human normal adrenal gland, combining single-nuclei RNA sequencing and spatial transcriptome data to reconstruct adrenal gland development and tumourigenesis. We identified two populations of potential progenitor cells resident within the adrenal cortex: adrenocortical progenitors NR2F2^+^-ID1^+^ cells, located within and underneath the capsule, and medullary progenitors SYT1^+^-CHGA^−^ cells, located in islets in the subcapsular region. Using pseudotime analyses, we provided evidence of the centripetal nature of adrenocortical cell development and of the essential role played by the Wnt/β-catenin pathway in the adrenocortical self-renewal. By comparing transcriptional profiles of cells of normal adrenal glands and adrenocortical adenomas we revealed a high heterogeneity with six adenoma-specific clusters. Overall, our results give insights into adrenal plasticity and mechanisms underlying adrenocortical tumourigenesis.

## Introduction

The human adrenal gland is a complex endocrine tissue that maintains homeostasis by responding to various physiological stimuli by secreting steroid hormones and catecholamines. The adult adrenal gland consists of two functionally distinct parts, the cortex and the medulla, which develop from different embryological origins^1^. According to textbooks, the adrenal cortex is organized in three zones bearing diverse morphological and functional characteristics. From the outer to innermost layer, histological analyses identify the zonae glomerulosa (ZG), fasciculata (ZF) and reticularis (ZR) involved in mineralocorticoid, glucocorticoid, and androgen synthesis and secretion, respectively. Beyond the cortex, the centre of the adrenal gland is occupied by catecholamine-secreting medulla, originating from neural crest cells.

Several animal model studies have been performed to reveal the architecture of adrenal glands and their development. According to these studies, the adrenocortical zonation is driven by several signalling pathways, including the Wnt/β-catenin, the hedgehog (HH), the cAMP/protein kinase A (PKA), and the insulin-like growth (IGF) pathways^2^. Among these, both Wnt/β-catenin and HH play an important role in the adrenal subcapsular area, where they presumably contribute to the centripetal renovation of the adrenal cortex^3^. The dysregulation of these signalling pathways is likely involved in human adrenal tumourigenesis^4^. However, having recent studies mostly been performed in mice, these findings cannot be directly transferred to humans.

Benign tumours deriving from the adrenal cortex (adrenocortical adenomas, ACA) are frequent in the general population (up to 10% in age > 70 years). They are usually incidentally discovered and mostly endocrine inactive adenomas (EIA)^5,6^. However, benign tumours may provoke autonomous steroid secretion associated with clinically relevant disorders, such as adrenal Cushing’s syndrome in cortisol-producing adenomas (CPA) or Conn’s syndrome in aldosterone-producing adenomas. On the other side, malignant adrenocortical carcinomas (ACC) are rare aggressive cancers with overall poor prognosis. In the last years, comprehensive genomics studies of benign and malignant tumours have identified alterations in signalling pathways involved in adrenocortical tumourigenesis (i.e.Wnt/β-catenin, Rb/p53 signalling and/or IGF system) and autonomous cortisol excess (i.e. cAMP/PKA pathway)^5–11^. However, the molecular mechanisms underlying the pathogenesis of a vast percentage of adrenocortical tumours remained largely unexplained. Therefore, a deeper understanding of the normal adrenal growth, differentiation and self-maintenance of adrenal cells in the adrenal tumourigenesis is urgently required.

Recently, single-cell/single-nuclei RNA-sequencing (sc/snRNA-seq) and spatial transcriptomics have provided unprecedented insights into cellular heterogeneity allowing the identification of in part unexpected cellular identities^12–14^. Particularly, cell atlases of normal human tissues based on scRNA-seq data may provide the basis for cellular analysis of pathological human samples - such as tumours - to gain a deeper understanding of tumourigenesis^15–17^. Whereas some recent scRNA-seq based studies focused on the neural compartment of the adrenal gland^18^, to our knowledge so far only one mouse study investigated on the single-cell transcriptome level the adrenal cortex by analysing the effect of stress on the hypothalamic-pituitary-adrenal axis ^19^. However, adrenal physiology significantly differs between rodents and humans.

Therefore, we aimed to provide a first comprehensive cell atlas of the cortex of the adult human normal adrenal gland (NAG) at single nuclei resolution including spatial information. This new approach allowed us to directly find new cell-specific human biomarkers and gain insights in molecular processes of adrenal self-renewal and homeostasis in humans. Furthermore, we integrated our atlas with snRNA-seq datasets from a clinical ACA cohort - comprised of EIA and CPA patient samples - to reveal human tumour-specific cell subpopulations and potential treatment targets.

## Results

### Single-nuclei transcriptome sequencing of NAGs

We sequenced 11,931 nuclei from six NAGs (Table 1 and Supplementary Fig. 1), obtaining an average depth of 25 million reads per sample. In an unsupervised cluster analysis, we identified six main cell clusters with distinct gene expression signatures. The identity of each cluster was assigned by cross-referencing upregulated transcripts with canonical markers from the literature. The dominant cluster was populated by classical adrenocortical cell types (Fig. 1A), including subclusters of ZG, ZF and ZR, as well as a new subcluster named “adreno-medullary progenitors” (AMP) (Fig. 1B). Highly expressed genes typical for these cortex subclusters are shown in Fig. 1C. The remaining “satellite” clusters were annotated as “Medulla”, “Fibroblasts & Connective tissue” (FC), “Stem/Progenitor and Vascular Endothelial cells” (Stem/P&VEC), “Myeloid cells” and “Lymphoid cells” (Fig. 1D). The top 100 differentially expressed genes (DEGs) defining the individual clusters of the NAG are listed in Supplementary Table 1.

**Fig 1.**
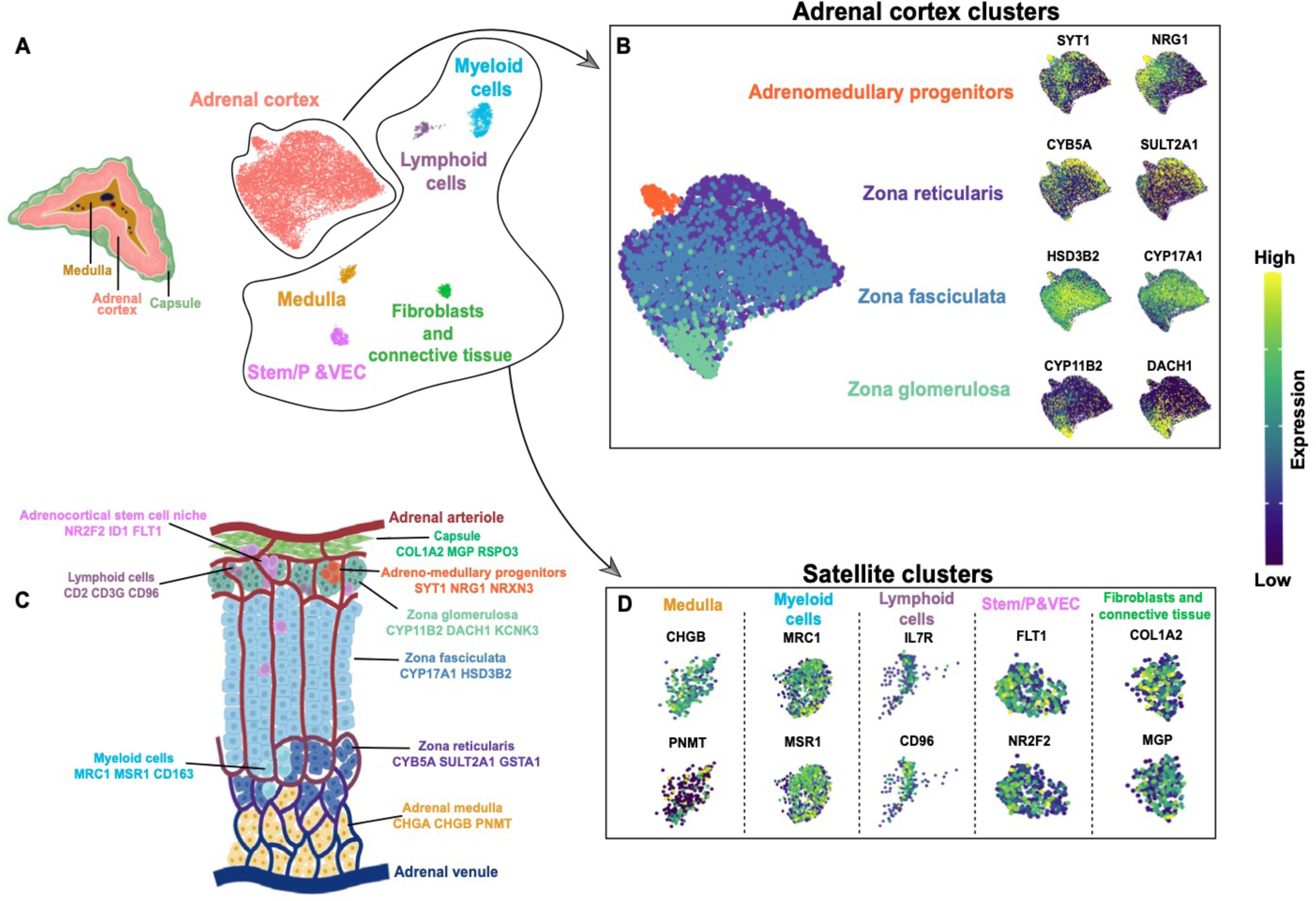
A comprehensive adult human normal adrenal gland cell atlas at single-nuclei resolution. **A.** Left: Transverse section depicting the three major adrenal zones (capsule, adrenal cortex, and medulla). Right: UMAP (Uniform Manifold Approximation and Projection) representation of the six integrated single-nuclei transcriptomic datasets (clusters were annotated by using known marker genes and DEG analysis). **B.** Adrenal cortex clusters: left: UMAP representation of the adrenal cortex clusters with complete zonation inferred by module scoring (see Fig. 2); right: feature expression plots representing genes specific for zonae reticularis, fasciculata, glomerulosa and adreno-medullary progenitors (scale represents log normalized expression values). **C**. Anatomical sketch of the adult human normal adrenal gland: the displayed markers for each identified cell type are based on DGE analysis in single-cell transcriptome data, available literature (see text for references) and in-situ validations by immunohistochemistry. **D.** Satellite clusters. Feature expression plots representing genes used in annotation of following clusters: “Medulla”, “Myeloid cells”, “Lymphoid cells”, “Stem/P&VEC“ (Stem/Progenitor & Vascular Endothelial cells) and “Fibroblasts & Connective tissue” (scale represents log normalized expression values).

### Zonation of the adrenal cortex

Within the adrenal cortex cluster, different cell subpopulations showed high expression of genes encoding for key steroidogenic enzymes representing the three cortex zones (Fig. 1B, Supplementary Fig. 2A-C, Supplementary Table 1). Cells from the ZG subcluster showed high expression of *CYP11B2*, as well as *DACH1* and *ANO4*, all previously described to be highly selective for this zone^20–23^. The ZF was characterized by high expression of *CYP17A1*, *CYP11B1*, and *HSD3B2*, whereas we observed elevated expression of *CYB5A*, *SULT2A1*, and *GSTA1* in the ZR^24,25^, as expected. Further gene enrichment analyses with DEGs from the three adrenocortical zones showed a significant overlap with regard to our annotation. In fact, aldosterone synthesis (fold enrichment (FE) >8), aldosterone-regulated sodium reabsorption (FE >6), endocrine and other factor-regulated calcium reabsorption (FE >4), and Wnt signalling (FE >2) were upregulated in the ZG subcluster, while cortisol synthesis and cholesterol metabolism were clearly overrepresented in the ZF (FE >23 and >13, respectively (Supplementary Fig. 3A-B). Steroid biosynthesis (FE >27), metabolism of xenobiotics by cytochrome P450 (FE >13), and estrogen signalling (FE >3) were higly represented in the ZR subcluster (Supplementary Fig. 3C). To further analyse the transcriptomic zonation of the adrenal cortex, we performed immunohistochemistry (IHC) to detect protein expression of selected mRNA-based markers in the different zones of the adrenal glands (represented via the kernel density^26^ in the “Uniform Manifold Approximation and Projection”, UMAP; Fig. 2A). Immuno-staining of DACH1 was uniform in the subcapsular region while CYP11B2 appeared in specific niches (Fig. 2C: DACH1, CYP11B2). These findings are consistent with the expression of the corresponding mRNAs in our single nuclei dataset: *CYP11B2* featured a low expression density in ZG but *DACH1* covered a substantial portion of this cluster (Fig. 2A: ZG). Moreover, CYP17A1 protein expression was detected in the entire cortex region (Fig. 2C: CYP17A1), corresponding to a higher RNA expression density in ZF with lower expression in the adjacent ZG and ZR clusters (Fig. 2A: ZF). In contrast, CYB5A protein staining clearly distinguished between ZF and ZR (Fig. 2C: CYB5A, SULT2A1). Although in the snRNA dataset both *CYB5A* and *SULT2A1* densities highlighted similar regions of the UMAP (Fig. 2: ZR), SULT2A1 was not selectively expressed in ZR as CYB5a, but was also expressed (at a lower intensity) in the ZG.

**Fig 2.**
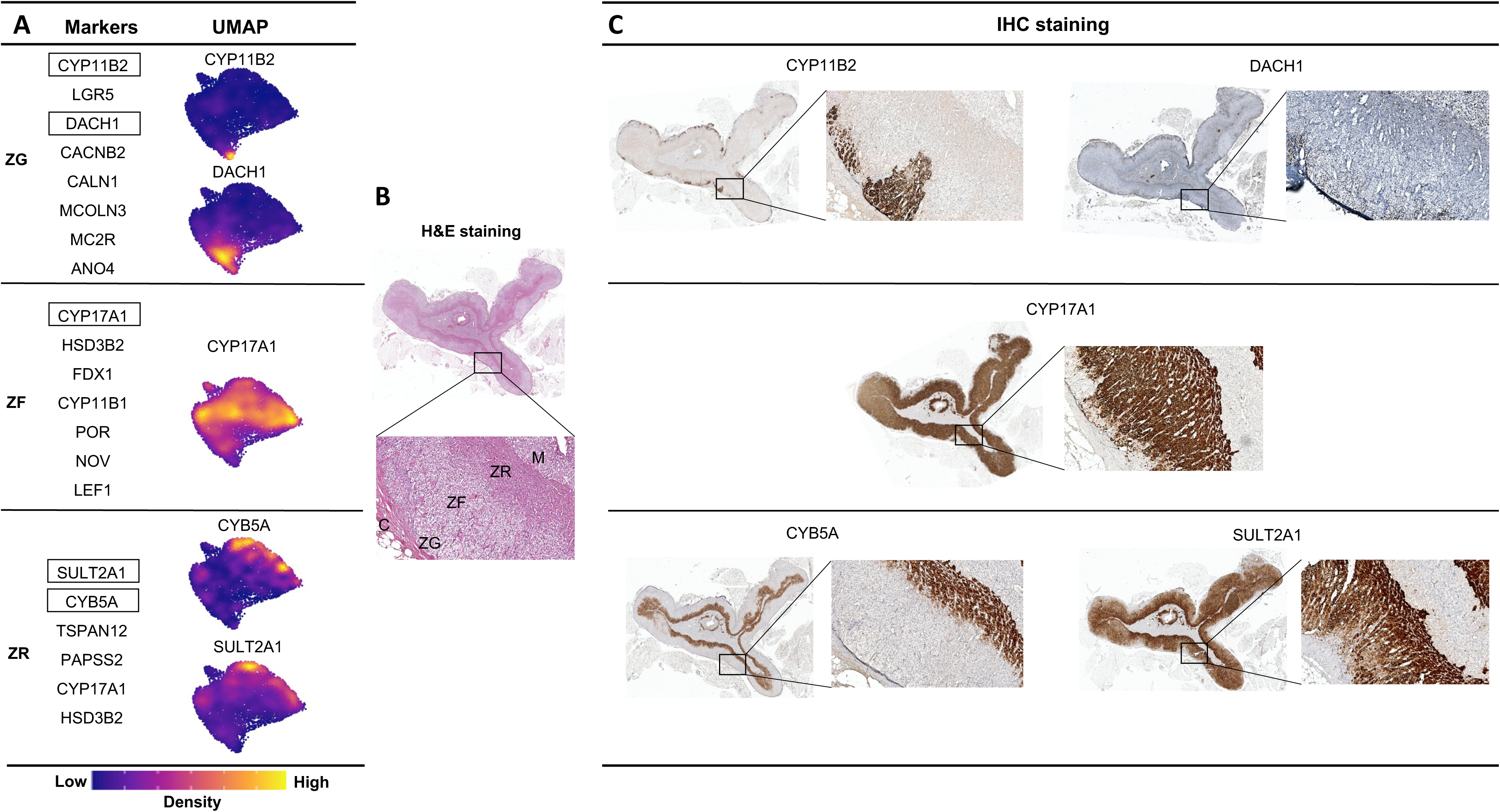
Correlation of single-nuclei transcriptome clusters with the zonation of the adrenal cortex. **A.** Kernel density estimation of selected features (in frames) for each of the three adrenocortical zones: ZG (*CYP11B2, DACH1*), ZF (*CYP17A1*) and ZR (*CYB5A, SULT2A1*). **B.** Example of hematoxylin and eosin (H&E) staining that served to distinguish the different zones of the normal adrenal gland (at 10x enlargement all the three zones of the cortex (glomerulosa, ZG; fasciculata, ZF; reticularis, ZR) together with the capsule (C) and the adrenal medulla (M) were observed). **C.** Consecutive adrenal sections were stained by immunohistochemistry (IHC) for the markers that are indicated in the boxes in A. CYP11B2 staining was confirmed to be expressed in the ZG as sporadic and scattered cells underneath the adrenal capsule or as clusters of cells (previously termed as aldosterone-producing cell clusters). DACH1 was expressed in the nuclei of the ZG cells. CYP17A1 was strongly expressed in the cytoplasm of adrenocortical cells of ZF and ZR, but not in the ZG. CYB5A1 was selectively stained in the cytoplasm of cells of the ZR, whereas SULT2A1 was stained in both ZF and ZR, although in the ZR a stronger staining was observed. All images were acquired by Leica Aperio Versa brightfield scanning microscope (Leica, Germany). 1x pictures with an enlargement of 10x of the same area of consecutive slides were used to obtain the IHC pictures.

### Adrenal progenitor and stem cells

Of note, within the main adrenal cortex cluster we identified a subcluster of sympathoadrenal lineage cells. These cells were characterised by transcriptional similarities to both neuronal cells and cells of the “Medulla” cluster (for comprehensive initial snRNA-seq analyses we included not only the cortex but also the medulla) (Fig. 1B, Supplementary Fig. 2F and Supplementary Table 1). As in the medulla, these cells specifically expressed genes involved in neurotransmission, such as *SYT1*, *NRG1*, *NRXN3*, *GRIK1*, and *CADPS*. Gene enrichment analysis further corroborated the chromaffin-like nature of these cells, revealing sets of genes involved in cholinergic and glutamatergic synapse (FE >7), glutamatergic synapse (FE >5), and axon guidance (FE >4) (Supplementary Fig. 3F). However, cells within this subcluster did not express markers of the mature medulla such as *CHGA* and *CHGB*, *TH*, *PNMT*, and *DBH*. Due to the expression of neuronal markers and the absence of hints to the mature medulla, we hypothesize that this subcluster represents early progenitors of the adrenal medulla, henceforth named as the “Adreno-Medullary Progenitors” (AMP) subcluster. We validated this hypothesis by using IHC to investigate the protein expression of SYT1 (synaptotagmin 1) and CHGA (chromogranin) (Fig. 3A-B). We observed low SYT1 expression in the entire cortex, whereas the medulla showed moderate expression. Interestingly, we found groups of cells with stronger SYT1 staining in the subcapsular region, which were organized as niches (Fig. 3B.1). In line with snRNA-seq data, these cells did not show CHGA expression in the double staining SYT1-CHGA (Fig. 3B.2), confirming that these niches may truly contain progenitor cells of the adrenal medulla. Interestingly, recent studies in murine models showed that nestin^+^ medullary progenitor cells were located not only in the medulla but also in niches of the mouse adrenal capsule, to contribute to the plasticity of the adrenal cortex. The here found presence of two different progenitor populations within the adult adrenal cortex might suggest a higher degree of flexibility of tissue renewal capacity than expected^27^ to ensure the maintenance of the adult adrenal cortex^28–32^.

**Fig. 3.**
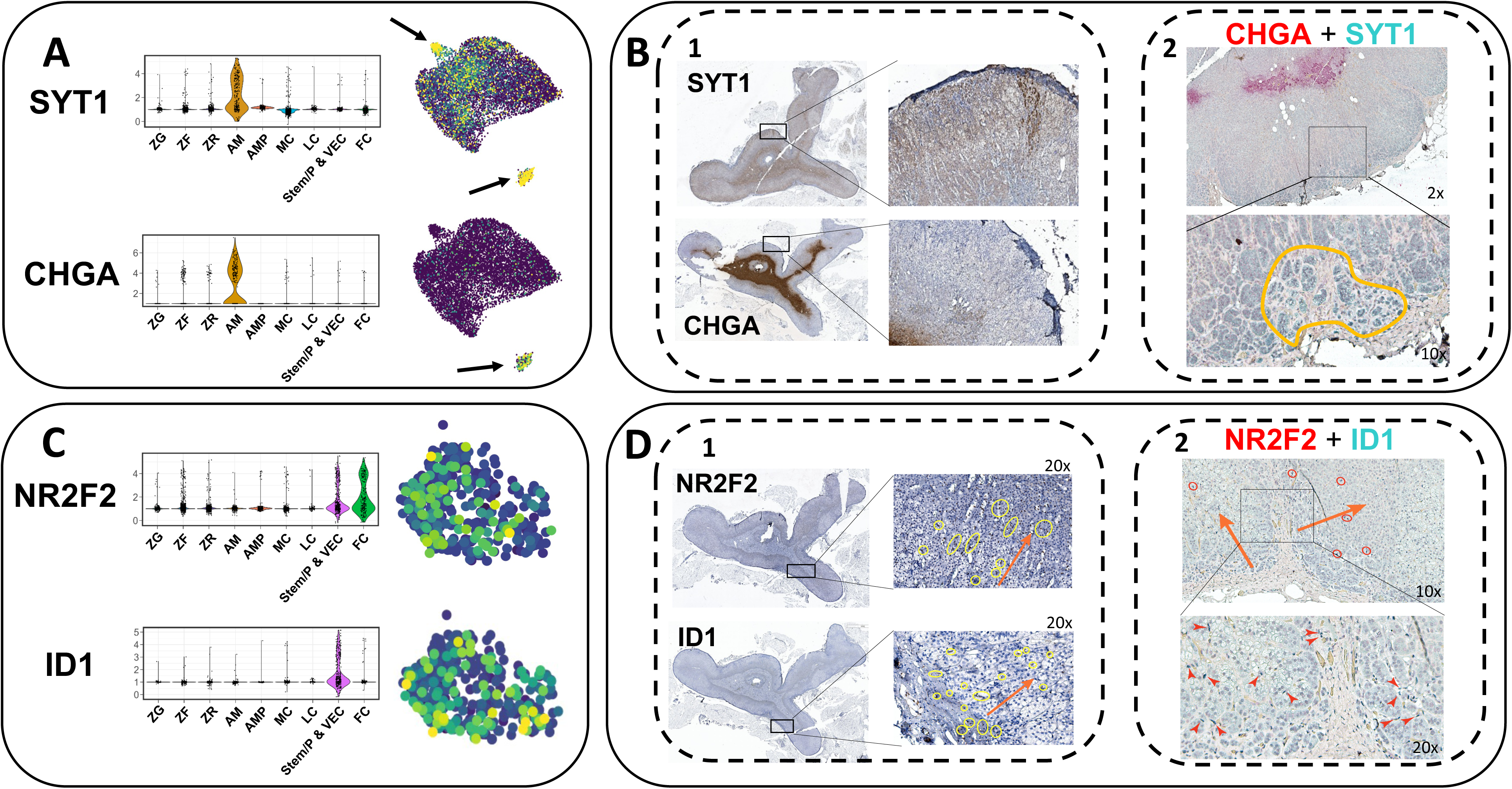
Adreno-medullary and adrenocortical progenitor cells in adult human normal adrenal glands. **A.** RNA expression of *SYT1* and *CHGA* in the AMP cluster. **B.1.** SYT1 and CHGA immunostaining. A low SYT1 protein expression was observed in cells of the entire cortex, whereas cells of the medulla showed moderate expression. Interestingly, groups of cells localized in the subcapsular region (organized as niches) showed a strong SYT1 staining. CHGA staining was found to be pronounced in medullary cells but was very low in cells of the inner part of the adrenal cortex and no staining was observed in the outer part of the adrenal cortex and in the adrenal capsule. **B.2.** CHGA/SYT1 double staining. At 2x enlargement, cells of the medulla showed a strong CHGA staining (fuchsin-red color). At 10x enlargement, a SYT1 strong positive niche (starker green-blue color) was marked with a yellow line. All images were acquired by Leica Aperio Versa brightfield scanning microscope (Leica, Germany). A 1x picture with an enlargement of 10x and 20x of the same area of consecutive slides were used to obtain the IHC pictures. **C.** Expression of *NR2F2* and *ID1* genes in the Stem/P&VEC cluster. **D.1.** NR2F2 and ID1 immunostaining. IHC analysis revealed sparse, rare cells mostly localized at the subcapsular levels that showed a high nucleus staining of NR2F2 or ID1 (marked with yellow circles). Those cells located under the capsule seemed to centripetally migrate from ZG to ZF and ZR (orange arrows represented the centripetal migration in adrenal glands). **D.2.** NR2F2/ID1 double staining. Nuclei of cells positive for both NR2F2 and ID1 were stained in dark blue (because of the sum of fuchsin-red + blue-green colours) and were mostly located under the capsule. Double positive NR2F2/ID1 nuclei were indicated with red circles at 10x enlargement and red arrows at 20x enlargement. Orange arrows at 10x enlargement represent the centripetal migration in adrenal glands.

Another cell cluster that includes a significant number of adrenal stem cell markers is the Stem/P&VEC cluster. Here, classical endothelial genes, such as *FLT1* (also known as *VEGFR1*), *KDR* (also known as *VEGFR2*), and *TEK*, as well as several genes associated with cell proliferation and differentiation, including *ETS1*, *ETS2*, *INSR*, and *DNASE1L3* were found (Supplementary Fig. 2E, Supplementary Table 1). Moreover, *NOTCH1*, a known marker of adrenal cortical progenitors^33^ (involved in cell fate specification, differentiation, proliferation, and survival) and its target gene *HES1* (responsible for the regulation of genome-wide glucocorticoid signalling^34^) were also both expressed at a high level (Supplementary Fig. 2E). Simultaneously, low gene expression levels of genes associated to steroidogenesis, i. e. *NR5A1* (also known as *SF1*) and *STAR* were found within this cluster. One more interesting finding was the distinct expression of *NR2F2* (also known as *COUP-TFII,* Fig. 1D), a potential marker of adrenocortical stem cells in mouse models^35,36^ and humans^37^. *NR2F2* expression was also observed in the “Fibroblasts and connective tissue” cluster (Fig. 3C) that shared a similar transcriptome profile to the adrenal capsule, expressing *COL1A2* and *RSPO3* (Supplementary Table 1). Recently, it was reported that *NR2F2* expression is most likely related to RSPO3^+^ capsular cells and that RSPO3 plays a direct role in the mouse adrenal homeostasis^28^. In addition to *NR2F2*, we found *ID1,* a known marker of mesenchymal stem cells, exclusively expressed in the same cluster. To gain more insight in the “stem” identity of this cluster, we performed IHC and microscopy-based tissue analysis with the undifferentiated protein markers NR2F2^35,36^ and ID1. NR2F2 staining was found in nuclei of rare, sparsely distributed cells. These cells were mostly localized at the subcapsular level, whereas few additional NR2F2^+^ cells were found within the inner cortex, following the trend of centripetal migration of adrenocortical progenitor cells from ZG to ZF to ZR (Fig. 3D.1). ID1 nuclei staining showed similar distribution when compared to NR2F2^+^ cells (Fig. 3D.1). Via double immunostaining, we found sparse NR2F2^+^-ID1^+^ cells mostly localized at capsular/subcapsular levels that appeared to migrate centripetally within the cortex, suggesting that ID1 may represent a new marker of stem/progenitor cells (Fig. 3D.2). Interestingly, a previous study also highlighted the role of ID1 in the tumourigenesis of adrenocortical cancer in dogs^38^. Pathway analysis corroborated our hypothesis, revealing significant enrichment of adherens junction (FE >5) and different signalling pathways, such as mitogen-activated protein kinase (MAPK, FE >3), Wnt pathway (FE > 2) and signalling regulating the pluripotency of stem cells (FE >2) (Supplementary Fig. 3E).

### Characterisation of additional adrenal cell clusters

As shown in Fig. 1, we detected four other cell types in the adrenal glands. The “Medulla” cluster showed an expression pattern typical for chromaffin cells with a predominance of *CHGA, CHGB, PNMT*, *DBH*, and *TH* (Fig. 1D, Supplementary Fig. 2F, Supplementary Table 1). Within the “Myeloid cells” cluster, we observed a large population of macrophages, mostly polarised into alternatively activated (M2) macrophages with high expression of *MRC1* (also known as *CD206*) and *CD163* together with *MSR1* (also known as *CD204*) (Fig. 1D, Supplementary Fig. 2G, Supplementary Table 1). Other highly expressed genes were *SRGN*, a proteoglycan that is typically expressed in myeloid and lymphoid cells, *CSF1R*, *CD86*, and *CD14* (Supplementary Fig. 2G). Furthermore, we observed a second immune cluster of lymphoid cells, which showed typical T cell markers, such as *IL7R* and *CD96* (Fig. 1D, Supplementary Table 1), as well as *CD247* (part of the T-cell receptor-CD3 complex), *RUNX3* and *ITK* (involved in T cell differentiation), *PRF1* and *GZMA* (indicators of granule mediated cytotoxic activity of natural killer and cytotoxic T cells) (Supplementary Table 1, Supplementary Fig. 2H). Of note, only few markers for B cells were found, e. g. *IKZF3* (involved in the regulation of B lymphocyte proliferation and differentiation) (Supplementary Fig. 2H and Supplementary Table 1).

The “Fibroblasts and connective tissue” cluster was characterised by cells that highly expressed collagen markers, such as *COL1A2* and *COL4A1*, and markers of the extracellular matrix, such as *MGP, FBLN1*, *LAMA2* and *CCN1* (Fig. 1D, Supplementary Fig. 2I and Supplementary Table 1). Using IHC, we confirmed protein expression of COL1A2 in the adrenal capsule and MGP in extracellular matrix and fibroblasts (Supplementary Fig. 4A).

Pathway analyses validated the satellite cluster annotations, revealing enrichment of cluster-specific gene sets, such as glutamatergic and cholinergic synapse (FE >8 and >5, respectively) for “Medulla”, phagosome and NF-kappa B signalling (FE >4 for both pathways) for “Myeloid cells”, Th1 and Th2 cell differentiation (FE >5) and B and T cell receptor signalling pathway (FE >4) for “Lymphoid cells” and focal adhesion (FE >5) and extracellular matrix-receptor interaction (FE >4) for “Fibroblasts and connective tissue” cluster (Supplementary Fig. 3F-I).

### Adrenal cortex zonation by spatial transcriptomics in NAG

We validated the cellular and tissue assigments of genes based on snRNA-seq data by spatial transcriptomics using the Visium assay in two consecutive adrenal gland sections deriving from the exemplary patient sample #NAG-7. As we further focused in this study on the adrenal cortex, the medulla was not part of the analysed tissue sections. To identify each zone, we transferred the cell type annotation from the microdroplet-based snRNA-seq into the Visium data set using a cell-to-spot correlation framework (Fig. 4). Mapping of the adrenal cortex zone markers confirmed the snRNA-seq dataset. ZG appeared as a thin line of cells towards the outer cortex, while the distribution of spots with high ZF score was rather ubiquitous. Moreover, spots with higher ZR scores seemed to locate closer towards the center of the tissue section, although a substantial overlap between ZF and ZR was observed (Fig. 4 and Supplementary Fig. 5). Regarding the satellite clusters, the label transfer predicted similar mapping for “Fibroblasts and connective tissue” and “Lymphoid cells” to the region surrounding the cortex, although the latter displayed a low overall mapping score, probably due to the low abundance of lymphoid cells (Fig. 4). Of note, “Fibroblasts and connective tissue” markers such as *COL1A2* and *RSPO3* were exclusively expressed in this region, while “Lymphoid cells” markers could not be found (Supplementary Fig. 5). The mapping of the “Myeloid cells” cluster showed a specific expression of immune markers mostly around the small blood vessels within the adrenal cortex (Fig. 4). Finally, the Stem/P&VEC mapped with a high prediction score in the outer cortex, specifically localized in a subcapsular niche, while a medium to low prediction score was observed in the inner cortex (Fig. 4).

**Fig. 4.**
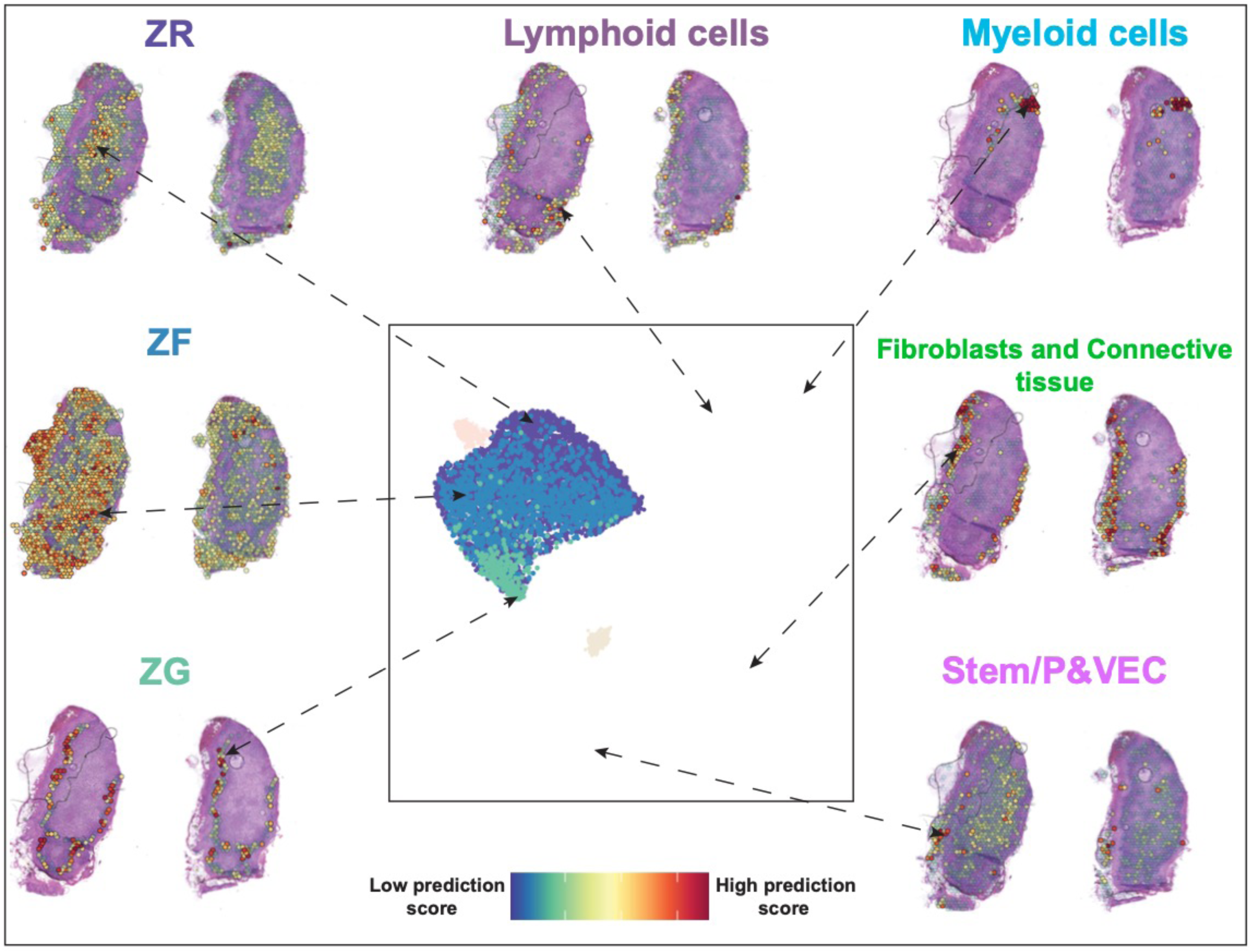
Validation of adrenal zonation by spatial transcriptomics in adult human normal adrenal glands. A pairwise (cell-to-spot) score is calculated within the label transfer framework in Seurat (V3) and projected onto sections. Two sections are represented for each mapped cluster. Medulla and AMP clusters were rendered transparent as the label transfer did not detect the presence of these cell types in the Visium assay.

### Pseudotime analysis confirms centripetal development of the adrenal cortex

Next, we aimed to understand the developmental relationship between the observed cell clusters within the adrenal cortex. To predict a differentiation trajectory, we performed pseudotime analysis^39^ using the spatial transcriptomics data. The integration of the two NAG replicates resulted in 5 distinct Louvain clusters (0 =Capsule, 1 =ZG, 2 =ZF-ZR, 3 =blood vessel, 4 =endothelial) that we annotated based on the results of the label transfer (Fig. 5A). As root node for pseudotime computation we selected cells expressing *RSPO3*, a well-established marker of capsule adrenocortical progenitors of mouse adrenal gland^23,28^ (within the capsule cluster, *RSPO3* was specifically expressed by a subset of cells (Fig. 5B)). The resulting unsupervised lineage trajectory of the adrenal cortex started with capsular cells, proceeded through the ZG cluster, and ended in the ZF-ZR cells, indicating a centripetal development of the adrenal gland.

**Fig. 5:**
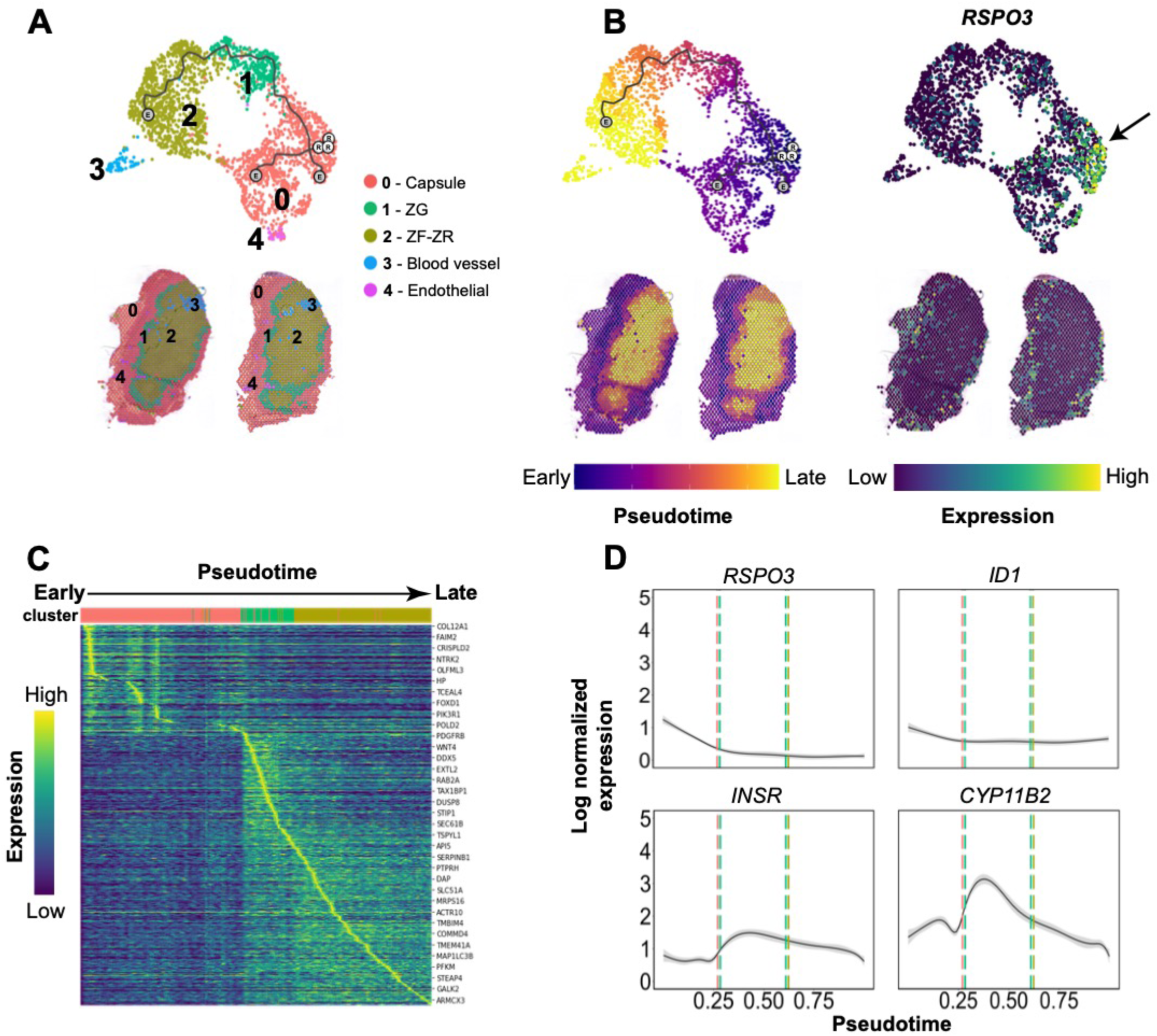
Trajectory analysis of spatial transcriptomics data in adult human normal adrenal glands. **A.** Top: Integrated UMAP (Uniform Manifold Approximation and Projection) from both tissue sections analysed using the Visium assay (10x Genomics). Trajectory was visualised as a black line. Bottom: clusters identified in the integrated object transferred to the tissue (Visium) sections. **B.** Pseudotime estimation both for the UMAP and the tissue sections: white circles represent the root node (chosen based on *RSPO3* expression, indicated with a black arrow (right panel)). **C.** Heatmap of 829 differentially expressed genes across pseudotime: cells were ordered by pseudotime (columns), genes were ordered by expression and pseudotime (rows). Top genes are highlighted. **D.** Expression variation of *RSPO3, ID1, INSR* and *CYP11B2* over pseudotime: smoothed lines were generated based on the scatter profile of each gene (95% confidence interval displayed in grey around the line). Vertical lines represent the transition zones from 0-Capsule (red) to 1-ZG (green) and 1-ZG to 2-ZF-ZR (ochre).

To further investigate potential changes in global gene expression dynamics along the trajectory, we applied pseudotime analysis in DEGs of the three main clusters (capsule, ZG, and ZF-ZR) (Fig. 5C). In total, we found 829 genes that significantly varied across pseudotime. Among them, *RSPO3* was comparatively strongly expressed in capsular, early pseudo-time cells, and weakly expressed within the inner-zones, late cell types (Fig. 5D). A similar pattern of cellular development was observed for *ID1*. *INSR* gene expression was specifically switched on in ZG (early time point in development) and its expression diminished throughout the inner cortex (later time points) (Fig. 5D). The selective activation of these genes at the early stage confirmed the stem/progenitor nature of the subpopulation of cells within the “Stem/P&VEC cluster” found in snRNA-seq analysis. Considering the steroidogenic-related genes, C*YP11B2* was first activated in the early ZG cells and significantly decreased throughout the ZG-ZR formation (Fig. 5D). We further observed an increased gene expression in the late ZG cells - and throughout the populations of the inner cortex - for *CXCR4* and *PRKAR1A* (cAMP/PKA pathway) and *ADGRV1* (cholesterol synthesis pathway) (Supplementary Fig. 6). Of note, the expression of *DLK1*, which is a proposed marker of a subcapsular cell population, was observed at two stages of the trajectory: a first peak appeared around the early development capsular region (Supplementary Fig 7A-B, indicated by a green box) and a second one at the ZG to ZF-ZR transition zone (Supplementary Fig. 7A-B, indicated by a blue box). Interestingly, *DLK1* expression visualized on the adrenal sections revealed a similar pattern described by Hadjidemetriou *et al.*^40^, i.e as “clusters of cells” (Supplementary Fig. 7B, regions inside blue boxes).

Moreover, the pseudotime analysis revealed interesting expression of Wnt/β-catenin pathway-related genes (Supplementary Fig. 6). *SFRP1*, *SFRP2* and *SFRP4* were found to be expressed in early or late capsular cells, but their expression decreased throughout the remaining cells subpopulations. *WNT4* was expressed in ZG only. Other genes such as *CTNNA1* or *CTNNA1L* were specifically switched on in the late ZG subpopulation and continued to be mildly expressed in the ZF-ZR zones (Supplementary Fig. 6).

### Integration of NAG and ACA snRNAseq data reveals adenomas-specific cells

Next, we used our human reference atlas to analyse cell type heterogeneity and identify possible tumourigenesis processes in our ACA patient samples (Table 2). For this, we sequenced 21,794 single-nuclei transcriptomes from 12 snap-frozen ACA samples (7 CPAs and 5 EIAs) and integrated this data with 5154 highest scoring NAG nuclei.

The differential expression analyses indicated that several genes involved in cell signalling, tissue remodelling, cell replication and RNA transcription were significantly upregulated in ACAs compared with NAGs (Fig. 6A.1-2). Particularly genes like *IGF2R* (average fold-change for gene expression, avg-log2FC >4.2, *p*<1e-6 in both EIA and CPA), *SLBP* (avg-log2FC >3.4, *p*<1e-6 in both EIA and CPA), *GAS2* (avg-log2FC >3.4, *p*=1.04e-272 in EIA and avg-log2FC >2.0, *p*=8.85e-153 in CPA), and *SP100* (avg-log2FC >1.8, *p*=2.45e-208 in EIA and avg-log2FC >1.6, *p*=9.33e-184 in CPA) were strongly expressed in both EIA and CPA (Fig. 6A.3). Genes like *MMP26* (avg-log2FC >3.6, *p*<1e-6) and *ADGRG2* (avg-log2FC > 1.5, *p*=2.27e-138) were strongly expressed in EIAs, whereas *RBFOX1* (avg-log2FC > 3.1, *p*<1e-6) and *HMGCS1* (avg-log2FC >1.6, *p*=4.51e-134) were predominantly expressed in CPAs (Fig. 6A.4).

**Figure 6.**
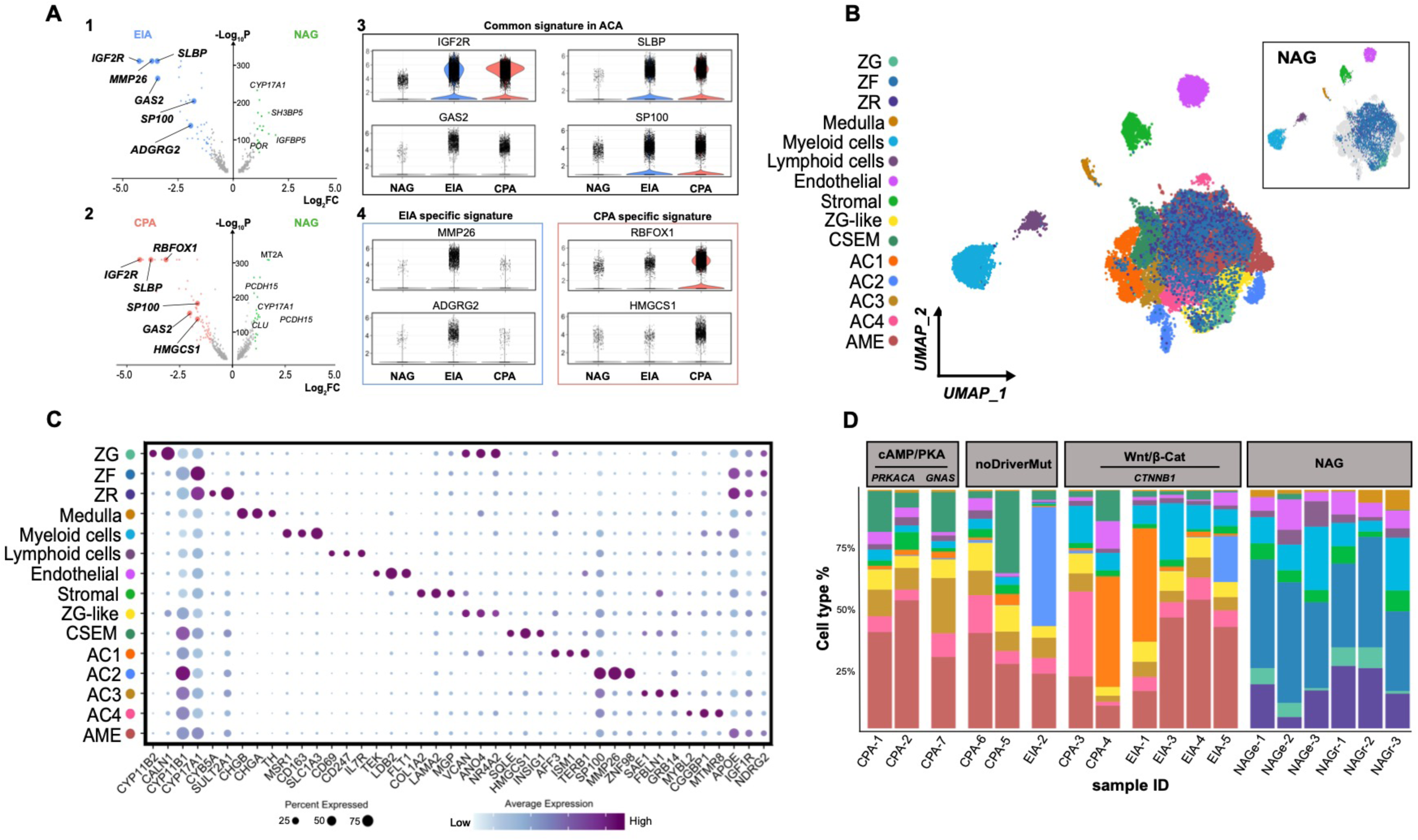
Identification of adenoma-specific clusters in adrenocortical adenoma (ACA) **A.1.** Volcano plot depicting differentially expressed genes between NAG and EIA. Genes characteristic of EIA subtype are written in bold. The x axis represents fold change (Log_2_FC) and y axis represents P values (-Log_10_P). **2.** Volcano plot depicting differentially expressed genes between NAG and CPA. Genes characteristic of CPA subtype are written in bold. **3.** Violin plots showing log normalized expression values of common signatures across the three subtypes (NAG, EIA, CPA). **4. Left:** Violin plots showing log normalized EIA specific gene expression values. **Right:** Violin plots showing log normalized CPA specific gene expression values. **B.** UMAP (Uniform Manifold Approximation and Projection) representation of the 18 integrated samples (normal adrenal glands (NAG, n=6); cortisol-producing adenomas (CPA, n=7); endocrine inactive adenomas (EIA, n=5) coloured by cluster identity. The UMAP could be divided into two groups: the “central cluster” containing previously identified cortex cell types (ZG, ZF and ZR) as well as ZG-like, Adenoma specific (AC1, AC2, AC3, and AC4), Cholesterol-and steroid-enriched metabolism (CSEM) and adenoma microenvironment (AME) clusters, and the “satellite clusters**”,** which are Medulla, Myeloid cells, Lymphoid cells, Endothelial and Stromal. NAG cell type distribution in the UMAP is represented within the top right box. **C.** Scaled average expression of selected markers for each cluster (dot size represents the percentage of cells in each cluster expressing the marker). **D.** Subtype and sample specific composition coloured by cluster identity: samples (x axis) are ordered by mutational signatures.

In line with the UMAP results of NAGs, in ACAs we found a central main cluster and 5 satellite clusters comprised of myeloid, lymphoid, endothelial, stromal (mainly composed of fibroblasts and connective tissue) and medullary cells (Fig. 6B). The central major cluster showed a complex organization including the three cortex-specific ZG, ZF and ZR and seven other subclusters (Fig. 6B-C). The “Cholesterol-and Steroid-Enriched Metabolism” (CSEM) subcluster was also found in a very small percentage in NAGs (~1%). The six remaining subclusters were specifically expressed in ACAs and were named as “Adenoma MicroEnvironment” (AME), “ZG-like”, and “adenoma-specific clusters” (AC1-4). These subclusters were mostly distributed in the periphery of the central major cluster, except for AME that occupied a large area in the central region (Fig. 6B). Top 100 DEGs defining the individual clusters of the integration of NAG and ACA are listed in Supplementary Table 2. In short, RNA markers specific for the ZF and ZR clusters of the NAG were largely expressed within the central cluster, and RNA markers such as *CYP17A1* and *CYP11B1* were rather ubiquitously expressed at low to medium levels across the tumour clusters (Fig. 6C), confirming the adrenocortical nature of these cells.

### ACA-specific clusters distribution according to subtype and mutational status

We further explored the cluster/cell type composition across the 18 samples by grouping our samples according to tissue entity (CPA, EIA or NAG) and mutational status – available from previous next-generation DNA or Sanger sequencing of bulk tumour tissues (no driver mutation, *CTNNB1* mutations, and *PRKACA/GNAS* mutations) (Fig. 6D).

The ZG-like and the AME clusters showed a homogeneous distribution, without significant differences among the samples (Fig. 6D). Within the “ZG-like” cluster, common ZG markers, such as *DACH1*, *ANO4*, and *NR4A2* (also known as *NURR1*) were dominantly expressed (Fig. 7A), indicating the presence of normal adrenocortical cells in ACAs. However, unlike ZG, the ZG-like cluster was characterised by lower to null expression of *CYP11B2* (avg-log2FC >-2, *p*=8.69e-36), *CALN1* (avg-log2FC >-2.3, *p*=9.60e-82) and *MC2R* (avg-log2FC >-1.9, *p*=1.17e-31). Moreover, ZG-like cluster showed also significant expression of non-ZG specific genes, such as *CYP11B1* (avg-log2FC >1.2, *p*=2.05e-16), *IGF2R* (avg-log2FC >4.4, *p*=3.79e-78) and *GAS2* (avg-log2FC >2.4, *p*=.82e-12). Pathway analyses revealed enrichment of several pathways involved in cell signalling, such as protein phosphorylation (FE >6), as well as pathways involved in calcium signalling (FE >3) and aldosterone secretion (FE >2) (Fig. 7B and Supplementary Fig. 8).

**Figure 7.**
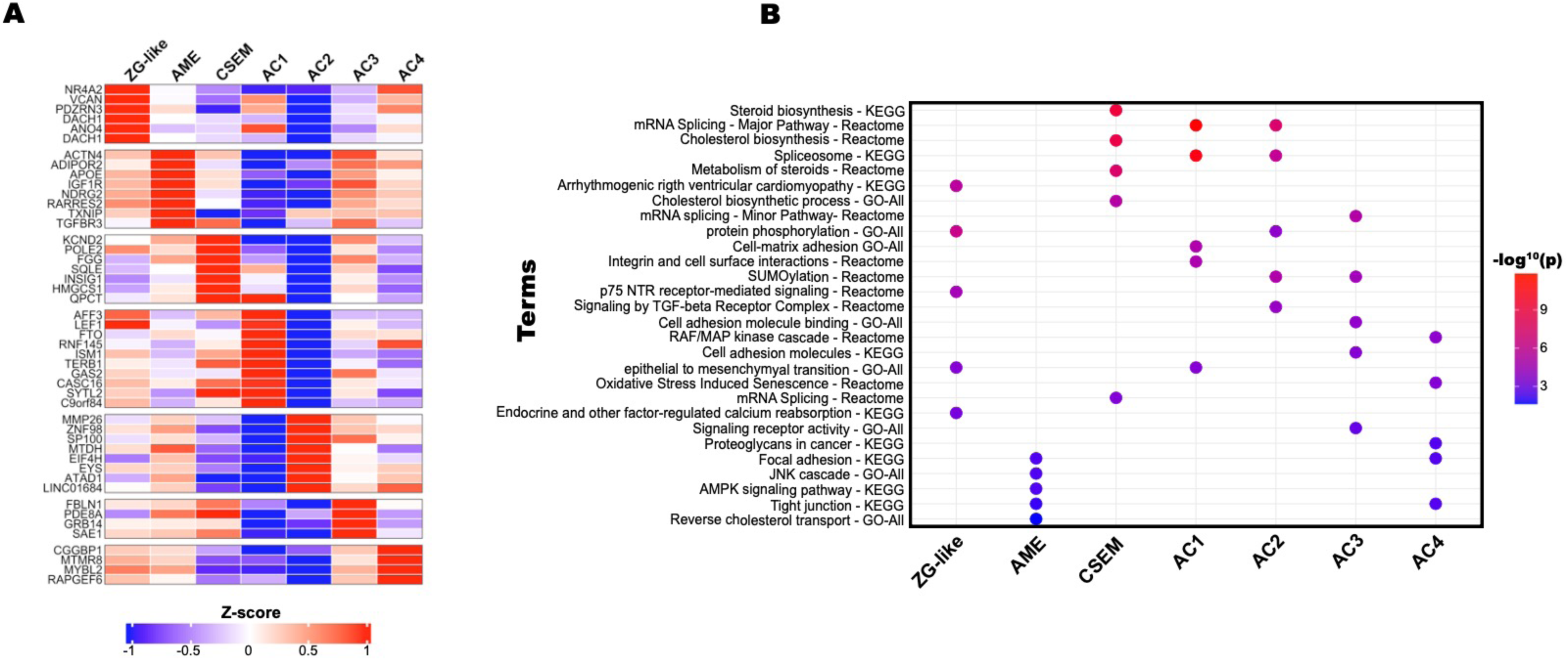
Transcriptomic profile of the newly identified ACA clusters. **A.** Heatmap representing top identified features for the 7 newly identified adenoma clusters. Colour scale represents Z-score computed from log normalized expression matrix. **B.** Top 5 significant pathways for each cluster combining results from 3 separate databases (KEGG, Reactome and GO-All (GO-MF, GO-BP, GO-CC))

Cells of the AME overexpressed genes like *APOE*, *ADIPOR2* and *RARRES2* involved in the lipoprotein’s metabolism, as well as *IGF1R* and *NDRG2* (Fig. 6C and 7A). *IGF1R* is a transmembrane tyrosine kinase receptor of the IGF family that activates a mitogenic signalling pathway, stimulating cell proliferation, division, and translation, and inhibiting apoptosis^41^. *NDRG2* regulates the Wnt signalling pathway by modulating *CTNNB1*-target genes^42^. Pathway analyses revealed enrichment of several pathways involved in cell signalling, such as AMPK (FE >4) and insulin signalling pathways (FE >3), cholesterol trafficking (FE >4), and pathways associated with different tight, focal and adherens junctions (FE >2) (Fig. 7B and Supplementary Fig. 8).

The CSEM cluster was significantly over-represented in the CPA cohort compared to EIAs (*p*=0.03; Fig.8A), with high expression of genes related to cholesterol metabolism, including *HMGCS1*, *SQLE*, and *INSIG1*, but also *KCND2*and *POLE2* (Fig. 7A), which are involved in DNA repair and replication. The enrichment pathway analyses confirmed this finding, showing several pathways involved in cholesterol metabolism and steroidogenesis, among which a very high enrichment of cholesterol and steroid biosynthesis (FE >9), pathways involved in ubiquitination processes, including protein ubiquitination (FE >5) and cellular response to oxidative stress (FE >2.5) (Fig. 7B and Supplementary Fig. 8).

**Figure 8.**
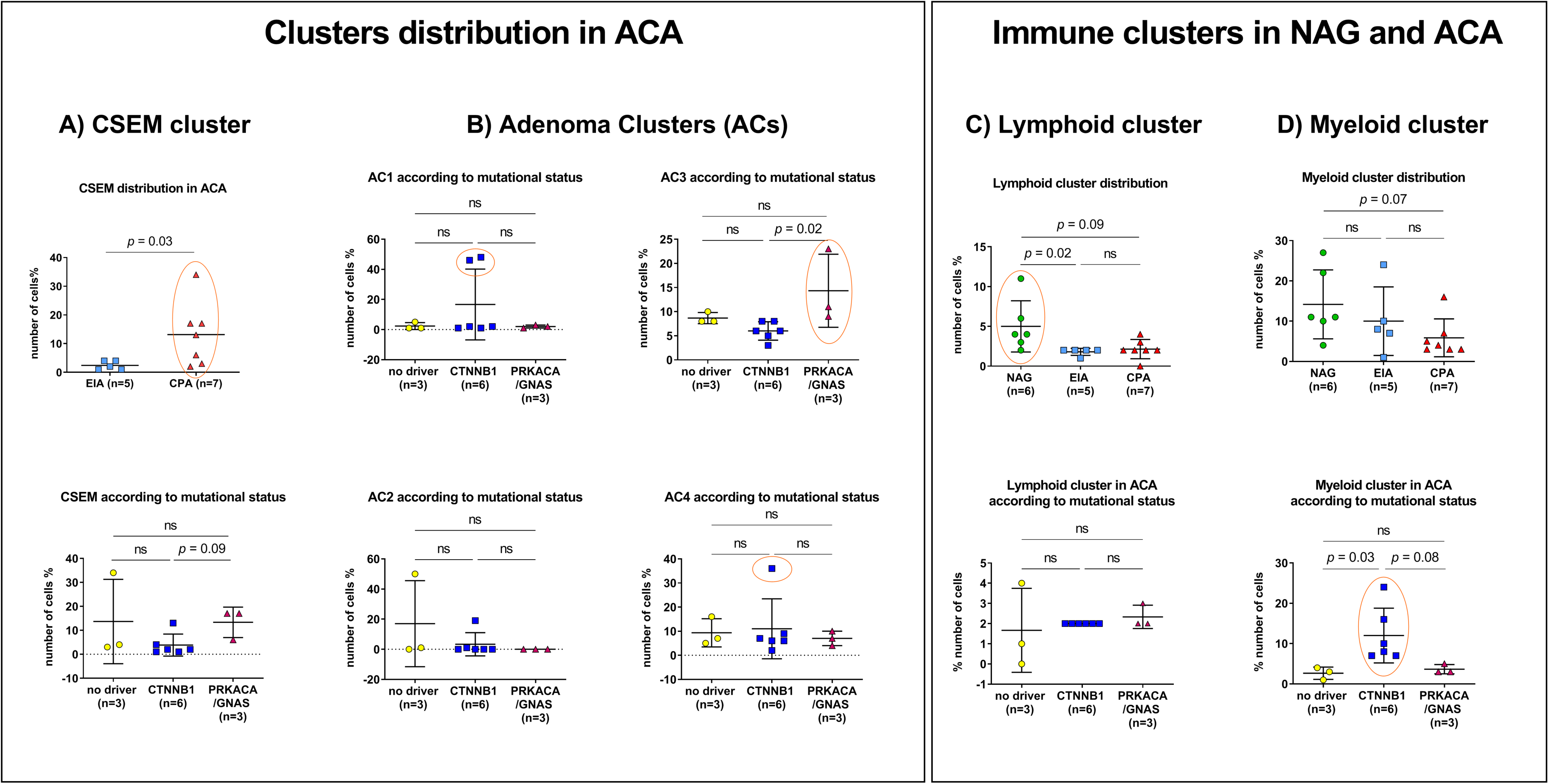
Cluster distribution in ACA samples across subtype and mutational status. Distribution of **A**. Cholesterol-and steroid-enriched metabolism (CSEM) cluster in cortisol producing adenomas (CPA) versus endocrine inactive adenomas (EIA; upper panel) and across mutational status (lower panel); **B.** Adenoma clusters (AC1-4) across mutational status; **C**. Lymphoid cluster across subtypes (upper panel) and mutational status (lower panel); **D**. Myeloid cluster across subtypes (upper panel) and mutational status (lower panel). Significant cluster enrichments are encircled in red.

The remaining clusters presented a slightly different distribution within the tumour subtypes and mutational status. Specifically, AC1 was mostly found in two *CTNNB1*-mutated samples, one EIA (EIA-1, ~48%) and one CPA (CPA-4, ~46%) (Fig 6D and Fig. 8B), with a high expression of the β-catenin target genes *LEF1*, *AFF3*, and *ISM1* (Fig 7A, Supplementary Table 2). Both *AFF3* and *ISM1* are known to be overexpressed in benign and malignant adrenocortical tumours with *CTNNB1* mutation^43–45^. Also, we found a high expression of *FTO*, a N6-methyladenosine that promotes cells proliferation, migration and chemo-radiotherapy resistance by targeting *CTNNB1* in different cancer types^46,47^ (Fig. 6C and Fig. 7A). In addition, other genes involved in cell proliferation, including *GAS2* and *TERB1*, and long non-coding RNAs, such as *CASC16* and *C9orf84,* were found to be highly expressed within this cluster (Fig. 7A). The enrichment analysis revealed a very high expression of different pathways involved in spliceosome (FE >10), cell proliferation and tissue development (FE >3), and extracellular cell-matrix adhesion (FE >5) (Fig. 7B and Supplementary Fig. 8).

AC2 cluster was more abundant in two EIA samples (EIA-2 and EIA-5) (Fig. 6D) without correlation with the mutational status (Fig. 8B). The AC2 cluster was characterised by an overexpression of genes like *MMP26*, *ZNF98* and *SP100* (Fig. 6C and 7A) associated with tumour promotion. Enrichment analysis showed also for AC2 a high expression of spliceosome pathway (FE >6) and ubiquitin-related pathways, including SUMOylation (FE >5) and cell cycle (FE >2) (Fig 7B and Supplementary Fig. 8).

In contrast, AC3 was significantly over-represented in CPAs with *PRKACA/GNAS* mutation compared to those with *CTNNB1-*mutated (*p*=0.02) (Fig. 6D and 8B). Particularly, it was very abundant in the single CPA with *GNAS* mutation (CPA-7, ~23%). Of note, we observed a high expression of *PDE8A* within this cluster (Fig. 7A). Also, a high expression of *SAE1*, associated to SUMOylation, *GRB14* and *FBLN1*, associated with cell growth and extracellular matrix, respectively, was found (Fig. 6C and 7A). Enrichment analyses confirmed a high enrichment in genes of the SUMOylation pathway (FE >4), cell adhesion molecular binding (FE >3), but also spliceosome (FE >2) (Fig 7B and Supplementary Fig. 8). Interestingly, AC1-3 showed a high enrichment of elements associated to spliceosome, sometimes differently expressed among the three. As such, splicing pathways were observed at very high levels in AC1 and decreased through AC2 and AC3, where they were found at lower levels.

Looking into the AC4 cluster, its distribution was similar among all tumour samples except for one CPA with *CTNNB1* mutation (CPA-3), where it represented ~36% of the cells (Fig. 8B). Within this cluster we found a high expression of the proto-oncogene *MYBL2*, as well as of *CGGBP1*, which regulates cell cycle in cancer cells (Fig. 6C and 7A). Pathway analyses showed significant enrichment of several pathways involved in extracellular matrix, cell-cell communication (FE >2) or different types of cells junctions (FE >2) and oxidative stress induced senescence (FE >3) (Fig. 7B and Supplementary Fig. 8).

### Immune clusters

We observed a decrease in quantity of both lymphoid and myeloid cells from NAGs to ACAs. Particularly, the lymphoid cluster was predominantly enriched in NAGs compared to EIAs (*p*=0.02) and by trend compared to CPAs (*p*=0.09) (Fig 8C). No difference was observed among the ACAs according to the mutational status. The expression of *UTY* significantly decreased from NAGs to both EIAs (avg-log2FC >-2, *p*=1.92e-08) and CPAs (avg-log2FC > 1.5, *p*=5.32e-09) (Supplementary Fig. 9A). Other genes like *HLA-A*, *HLA-B*, *HLA-C*, and *CCL5* followed a trend to decreased expression in CPA compared to NAG (avg-log2FC >-1.3, *p*=2.84e-08 for *HLA-A*, avg-log2FC >-1.2, *p*=6.55e-07 for *HLA-B*, avg-log2FC >-1.7, *p*=5.58e-07 for *HLA-C*, avg-log2FC >-1.5, *p*=1.05e-06 for *CCL5*) (Supplementary Fig. 9A).

The myeloid cluster was similarly represented in NAGs when compared to EIAs, but less represented in CPA samples (*p*=0.07) (Fig. 8D). Interestingly, the distribution of the myeloid cluster among the ACA samples changed according to the mutational status. Particularly, ACA samples with *CTNNB1* mutation revealed a higher abundance of myeloid cells compared to samples with no driver (*p*=0.03) and *PRKACA/GNAS* mutations (*p*=0.08) (Fig. 8D). Among differentially expressed genes (Supplementary Fig. 9B), *MSR1* was significantly downregulated in EIAs (avg-log2FC >-1.2, *p*=2.5e-19) and CPAs (avg-log2FC =-0.9, *p*=4.49e-12) compared to NAGs. Similarly, *SRGN* decreased in EIA (avg-log2FC >-1.2, *p*=5.24e-27) and CPA (avg-log2FC >-0.6, *p*=3.57e-11) compared to NAGs. Genes like *CSF1R* and *CD74* were less expressed in CPAs compared to both NAGs (avg-log2FC >-0.9, *p*=2.07e-07 for *CSF1R* and avg-log2FC - 0.9, *p*=1.87e-10 for *CD74*) and EIA (avg-log2FC >-1.0, *p*=3.04e-10 for *CSF1R* and avg-log2FC >-1.3, *p*=4.38e-24 for *CD74*) (Supplementary Fig. 9B).

## Discussion

In this study, we provide a high-resolution, comprehensive transcriptional cell atlas of the normal adult human adrenal gland including spatial RNA and protein expression information. We could distinguish characteristic zones of the adrenal cortex (ZG, ZF, ZR), as well as the adrenal medulla and capsule. In addition, we found different satellite clusters, including lymphoid, myeloid and vascular cells. Moreover, by comparing our single-cell transcriptome atlas of the adult human normal adrenal glands to a cohort of adrenocortical adenomas, we revealed seven adenoma-specific cell types that were involved in adrenocortical tumourigenesis.

One of the key findings of our study was the identification of two different populations of potential progenitor cells within the adult human NAG: the adrenocortical progenitor-like NR2F2^+^-ID1^+^ cells, located within and underneath the capsule, and the medullary progenitor-like SYT1^+^-CHGA^−^ cells, gathering in niches in the subcapsular region of the adrenal cortex. To our knowledge, we here show for the first time the presence of NR2F2^+^ progenitor cells in adult human adrenal glands. Moreover, we also found the co-expression of NR2F2 with ID1 (a mesenchymal stem cell marker) in few cells that were predominantly located under the capsule. As observed for progenitor cells in animal models, also in human adrenal glands these subcapsular cells showed a centripetal migratory pattern. The high differentiation potential of ID1^+^ cells is supported by high expression within the same “Stem/P&VEC cluster” of genes associated with cell proliferation and differentiation (*ETS1*, *INSR*, *HES1*, *DNASE1L3*, *NOTCH1)*^33^. Moreover, the trajectory analysis of spatial transcriptomic data enabled us to observe the selective activation of some genes at the early stages of pseudotime, confirming the progenitor nature of this subpopulation. In addition, the microenviroment of this cluster is rich in adherens junctions and vascular cells, which are essential in cell-cell communication, cell renewal and differentiation^48,49^. All these findings suggest that these ID1^+^ cells might represent a new progenitor population in the adult adrenal cortex.

In addition, we observed a small subcluster of cells, which expressed neuronal-like gene markers, including *SYT1*, *NRG1*, *NRXN3*, *GRIK1* and *CADPS* as found in chromaffin cells of the adrenal medulla. The subcluster was void of markers of mature medullary cells such as *CHGA*. Using IHC we validated the presence of SYT1^+^-CHGA^−^ cells that form clusters in the subcapsular region of the cortex, as had been observed for the nestin^+^ medullary progenitor cells in mouse. However, we could not identify nestin itself, potentially due to the differences in gene expression between mice and humans. Nevertheless, we cannot exclude that the rarity of these progenitors in adult adrenal glands or their presence only in early-stage adrenal development is responsible for this fact. The neuronal-like nature of the expressed markers within this subcluster raised the question if SYT1^+^-CHGA^−^ cells could represent a progenitor population of the adrenal medulla that is located in the subcapsular region of the adult human adrenal cortex, as had been observed in animal models^50^. Moreover, in line with the undifferentiated nature of NR2F2^+^-ID1^+^ and SYT1^+^-CHGA^−^ cells, both potential progenitor clusters showed little to no expression (< 5% in both clusters) of mature cells markers at transcriptional levels, including *NR5A1* and *STAR* in the Stem/P & VEC cluster, and *CHGA*, *CHGB*, *TH*, *PNMT*, and *DBH* in the AMP cluster.

Interestingly, by combining pseudotime and spatial transcriptomic analysis, we corroborated the centripetal nature of adrenocortical cell development. However, it still remains to be demonstrated ex vivo whether NR2F2^+^-ID1^+^ and SYT1^+^-CHGA^−^ cells represent populations capable to undergo steroidogenic or chromaffin differentiation. Future lineage tracing experiments will enable us to functionally characterise these progenitor subpopulations, this being beyond the scope of this pilot study.

The trajectory and pseudotime analyses allowed us to look more deeply into the transition zone between the capsular and sub-capsular region, where the progenitor cells are mostly located. Here we showed, for the first time, a selective activation of members of the Wnt/β-catenin pathway (i.e. *SFRP2* and *SFRP4*), which play an important role in adrenocortical renewal and differentiation^51^, as well as in adrenocortical tumourigenesis^11,52,53^. Particularly, both *SFRP2* and *SFRP4* were expressed in human mesenchymal stem cells isolated from bone marrow^54^, and a decreased expression of *SFRP2* was associated with aldosterone-producing adenoma development^55^. These findings corroborated an essential role of the Wnt/β-catenin pathway for progenitor cell formation and function throughout the adrenal development. Intriguingly, *DLK1* expression was observed in cells organised as clusters in the subcapsular region, as previously described^40^. Pseudotime analyses revealed that *DLK1* expression exhibits two peaks (first during the capsule-ZG transition and second during ZG to ZF-ZR transition), strengthening its potential role in adrenocortical remodelling.

Recently, it was proposed that adrenal renewal is sexually dimorphic^36,56^. In our study, we were not able to find any significant transcriptomic differences between the two sexes, potentially due to the small sample size (tissue from two females versus four males) and the limited number of (few hundreds of) detected expressed genes per nucleus. Similarly, age-specific differences were not observed, as all samples of this cohort belonged to adult subjects (ranging from 49 to 81 years of age).

Our transcriptome atlases of healthy adult human adrenal glands will be an important tool as reference to investigate adrenal-related disease. In a first step, we interrogated benign adrenocortical tumours (adenomas), which were further classified according to glucocorticoid secretion and mutational status. Using this approach, a high intratumoural heterogeneity was observed, confirming for the first time at single-nuclei level previous morphological and genetic findings in bulk tumours^57–59^.

The comparative analysis of the transcriptional profiles of NAGs and ACAs revealed the presence of seven adenoma-specific clusters. First, our results revealed an adenoma microenvironment (AME) cluster, which was quite homogeneously distributed among the samples. This represents a lipid-enriched milieu, similar to the tumour microenvironment described in other solid tumours^60^. It has been demonstrated that such an enrichment in adipocyte and lipid metabolism related genes plays a key role in aberrant cell growth and cancer progression, as well as in immune escape and treatment-resistance^60^. Furthermore, as reported for the tumour microenvironment in colon cancer^61^, a high expression of *RARRES2*, a key modulator of tumour growth and inflammation, was detected within this cluster. This finding is in line with a previous study reporting a higher *RARRES2* expression in benign compared to malignant adrenocortical tumours^62^. Within the AME, also a high expression of *IGF1R*, which plays an important role in the adrenocortical tumourigenesis, was observed^41^. Interestingly, an overepression of *IGF1R* within the tumour microenvironment was reported to promote tumour initiation and progression in lung cancer and medulloblastoma^63,64^.

The “cholesterol- and steroid-enriched metabolism” cluster, which showed a high expression of cholesterol pathway genes (i. e. *HMGCR*, *MSMO1*, *INSIG1*, *HMGCS1*), was, as expected, largely represented in CPAs^65^. Furthermore, in the single CPA with *GNAS* mutation (CPA-7), we observed a high prevalence of AC3, characterised by an overexpression of *PDE8A*. These results confirm previous reports that highlighted the potential role of the phosphodiesterase 8 family members in the pathogenesis of Cushing’s syndrome^11,66^. On the contrary, the AC2 cluster was significantly abundant in two out five EIA samples (EIA-2 and EIA-5), where a high enrichment of post transcriptional modifications (SUMOylation, deubiquitination) was detected.

A high morphological heterogeneity was particularly observed in *CTNNB1*-mutated adenomas, as recently reported in hepatocellular carcinomas harbouring mutations in the β-catenin gene^67,68^. Based on the transcriptome profile, we observed three types of *CTNNB1*-mutated ACA: one with considerable abundance of AC1 (more than 45% of cells, including one EIA and one CPA); one CPA with abundance of AC4 (more than 35% of cells); and the remaining three EIAs without preference for a specific adenoma cluster. Of note, in the CPA with abundance of AC1 we have observed a high expression of *AFF3*, *ISM1* and *LEF1*, all known *CTNNB1* target genes in benign and malignant adrenocortical tumours^43–45^. Particularly, it has been demonstrated that *AFF3* mediates the oncogenic effects of β-catenin in ACC cell line NCI-H295R by acting on transcription and RNA splicing^44^. Similarly, in the AC4 abundant CPA, a high expression of *MYBL2* was found. It has been shown that *MYBL2* is a Wnt/β-catenin target gene in different types of cancer-xenograft models^69^. Furthermore, *MYBL2* regulates cell proliferation, survival and differentiation and its overexpression is associated with poor outcome in different cancers types^70^, including ACC^71^. These findings might support the previously reported hypothesis of a potential adenoma-carcinoma sequence at least in a subgroup of *CTNNB1*-mutated adrenocortical tumours^11^.

Another notable finding emerging from the transcriptional profiles of adenomas was the enrichment of elements associated with spliceosome observed within three ACs (AC1-3), suggesting a potential role of spliceosome pathway in adrenocortical tumourigenesis, as demonstrated in other benign and malignant tumours^72–74^.

Interestingly, a different distribution of immune clusters emerged among tissue entities. Both lymphoid and myeloid clusters were better represented in normal adrenals than in adenomas. Although this trend was more evident for lymphoid cells, the median number of myeloid cells showed a tendency to decrease in CPA compared to EIA and NAG. This finding is in line with the known interplay between glucocorticoids and tumour-immune infiltration observed in malignant adrenocortical tumours^75^, suggesting the inhibition of tumour-infiltrating immune cells also in CPAs. In addition, we demonstrated for the first time a high prevalence of M2-like macrophages within the myeloid cluster in normal adrenal. M2-like macrophages have anti-inflammatory properties and promote cell homeostasis, confirming the potential trophic function of adrenal macrophages^76^. This shifted macrophage polarisation towards M2-like phenotype and, in particular, towards M2c subgroup, characterised by a high expression of the scavenger receptors *MRC1* and *CD163*^77^, could be related to the autocrine action of glucocorticoids secreted by the adrenal gland. To note, in adenomas, the mutational status significantly affected the myeloid cluster, which was highly enriched in *CTNNB1*-mutated adenomas. As observed in NAG, adenoma-associated macrophages presented a prominent M2-like polarisation, corroborating previous studies reporting that Wnt/β-catenin signalling promotes M2-like macrophage in tumours^78^ and that the M2 signature is highly prevalent in *CTNNB1*-mutated samples^79^.

### Short summary and outlook

In conclusion, using snRNA-seq and spatial RNA and protein expression analyses of adult human adrenal glands, we provide a first comprehensive cell atlas to identify complex mechanisms of homeostasis and self-renewal of the normal adrenal gland. Moreover, the present reference atlas allowed us already to analyse molecular processes of adrenocortical tumourigenesis and autonomous cortisol secretion in adenomas. In the future, further improved snRNA-seq technologies that will generate significantly more sequences per nucleus and provide data from more nuclei per analysis will help to ameliorate transcriptional data analysis, for example to deeper characterise developmental processes of the two newly identified progenitor cell types (SYT1^+^-CHGA^−^- and NR2F2^+^-ID1^+^-cells) in NAGs. Furthermore, ex vivo experiments will also be needed to analyse in more detail the influence of markers such as DLK1 and ID1 on cellular differentiation and specification.

## Materials and Methods

### Tissue sample selection

For single-nuclei RNA-seq (snRNA-seq), nuclei were isolated from six snap-frozen samples collected from NAGs deriving from the tissue surrounding EIA or from adrenalectomies performed during surgery for renal cell carcinoma (Supplementary Fig. 1). Demographic data and details of detected cellular transcriptomes are summarized in Table 1. Additionally, single nuclei were isolated from 12 fresh-frozen ACA samples, including 7 CPAs and 5 EIAs. Among these samples, 4 tissues were included in two previous studies on whole-exome sequencing^10^ or bulk RNA-sequencing^11^. Clinical characteristics, hormonal secretion, and detected cellular transcriptomes are summarized in Table 2 and Supplementary Table 3. For validation by immunohistochemistry (see below), 16 formalin-fixed paraffin-embedded (FFPE) tissue sections of NAGs were analysed (Supplementary Table 4). FFPE tissues included 3 out of 6 NAGs evaluated by snRNA-seq and one sample used for the Visium analysis.

Tissue samples were selected from the Würzburg Adrenal Biomaterial Archive, part of the BMBF-funded Interdisciplinary Bank of Biomaterials and Data Würzburg (IBDW) applying the highest standards for biobanking^80^ (details in Supplementary Material).

The study was approved by the ethics committee of the University of Würzburg (No. 93/02 and 88/11) and written informed consent was obtained from all subjects.

### Sequencing of driver mutations in adenoma tissues

Known driver hot-spot mutations in *CTNNB1* (exon 3), *PRKACA* (p.Leu206) and *GNAS* (p.Arg201 and p.Gln227)^10,11^ were evaluated by Sanger sequencing in bulk tumour tissues (n=8). Four samples were analysed by Whole Exome Sequencing or RNA-seq as part of previous studies from our group^10,11^. DNA was isolated from snap-frozen ACA tissues with the Maxwell® 16 Tissue DNA Purification Kit (#AS1030, Promega, Madison, WI, USA) according to the manufacturer’s instructions. The mutations were genotyped by Polymerase Chain Reaction (PCR), as previously reported^81^ (details in Supplementary Material).

### Clinical data collection

For ACAs, the endocrine workup was conducted according to the adrenal incidentalomas guidelines issued by the European Society of Endocrinology/European Network for the Study of Adrenal Tumors^5^ and the Endocrine Society guidelines for the diagnosis of Cushing’s syndrome^82^. Hormone levels were measured using commercially available analytical procedures (details in Supplementary Material), as previously reported^83^. CPA derived from patients with overt Cushing’s syndrome and unequivocal laboratory results (for details see Supplementary Material).

### Single-nuclei RNA sequencing (snRNA-Seq)

#### Single-nuclei isolation

We applied a previously published protocol for nuclei isolation from snap-frozen tissue^84^ (Supplementary Fig. 1 and Supplementary Material). The nuclei integrity and purity were checked under a microscope and the yield was quantified using a Neubauer chamber. ***Single-nuclei RNA sequencing:*** We used the inDrop™ method consisting of three major steps: Silanization, Reverse Transcription and Library preparation (1CellBio™, USA) following manufacturer’s protocols. First, chips were silanized, primed with the inDrop hydrogel beads, droplet making oil, RT/lysis mix and the nuclei. After completing the run protocol, cell encapsulations were collected and exposed to UV light to release photocleavable primers from hydrogel beads. Subsequently, reverse transcription was performed, and the beads filtered out. From there on, the library preparation step followed the CEL-Seq 2 protocol^85^: the RT product was digested by ExoI and HinFI and purified using AMpure XP beads (Beckman Coulter™, USA). Then, the second strand cDNA synthesis and *in vitro* transcription (IVT) steps were carried out. The resulting RNA was subjected to fragmentation and then reverse-transcribed using random hexamers. The cDNA was ultimately purified using AMpure XP beads and quantified by qPCR assay^86^ (Roche Light Cycler 480 Instrument II, Switzerland).

After quantification, the 6 normal adrenal gland (NAG), 2 endocrine inactive adenoma (EIA-3, EIA-4) and 2 cortisol producing adenoma (CPA-1, CPA-2) libraries were pooled together and sequenced using an Illumina HiSeq 2000 instrument with 60 bases for read (R1), 6 for the Illumina index and 50 for the read 2 (R2). The 6 samples were integrated using the standard Seurat (version 3.0) integration pipeline^87^ (freely available software): The resulting data is shown in Supplementary Fig. 1C. Cells from the two subtypes showed a similar clustering pattern, suggesting that the data were well integrated. Of note, data derived from either NAG-EIA or NAG-RCC patients showed correlation with a high Pearson correlation score of 0.99 (Supplementary Fig. 1D), indicating that NAGs could be merged in one single group. Moreover, the remaining 8 adrenocortical adenoma (ACA) libraries were pooled together and sequenced on two NovaSeq S1 flow cell at 2 x 100 configuration.

#### Generation of count matrices and core processing

The resulting base calls (BCL) files were converted to FASTQ files using bcl2fastq tool (Illumina). These raw sequencing reads were processed with the zUMI^88^ (version 2.5 – default parameters) pipeline. The count matrices from reads spanning both introns and exons (inex) were used for subsequent analysis. Prior to pre-processing, the genes/features were converted from Ensembl (ENSG) IDs to gene symbols using a custom script. The same cut-off was used to filter nuclei and genes likewise from individual samples (min.features= 200 and min.cells =3, *CreateSeuratObject*). The fraction of reads mapping to the mitochondrial chromosome (MT-) was calculated (*PercentageFeatureSet*).

#### Data normalization and integration using Seurat

The filtered data was log-normalized (*NormalizeData*) and 2000 highly variable genes (HVG) were selected. The six samples were then integrated (1. *FindIntegrationAnchors, 2. IntegratedData*)^89^. The integrated data was then scaled (*ScaleData)* and the effect of mitochondrial genes was regressed out.

#### Dimensionality Reduction and Clustering

Thirty principal components (PCs) were computed (*RunPCA*) and first 15 were used to generate the UMAP reduction (*RunUMAP*). Subsequently, the k-nearest neighbours (knn) were determined (*FindNeighbors*) and clustering was performed (*resolution=0.01, FindClusters)*. The resulting clusters were annotated using known marker genes and differential gene expression (DGE – *FindAllMarkers*) analysis.

#### Module Scoring

A list of top DEGs and prior knowledge was curated for each cluster, and combined gene expression scores were calculated (*AddModuleScore*). The Seurat object was further filtered using these scores, with the following thresholds: ZG >0.3, ZF >0.8, ZR >0.5, MC >0, LC >0, FC >0, Stem/P&VEC >0, AM >0, AMP >0. This resulted in a count matrix of 5154 nuclei that was then integrated with the ACA dataset. A Shiny implementation (R) was set up to easily navigate within this single-nuclei atlas.

#### Gene enrichment analysis

We used pathfindR^90^ to perform the gene enrichment analysis using the *FindAllMarkers* output table from Seurat. The analysis was performed using the function *run_pathfindR* for both NAG and ACA samples. For the NAGs, KEGG pathways was used as reference. To better cover tumour heterogeneity in ACAs, three different databases (KEGG, Reactome and GO-All) were used as reference. For each ACA cluster, similar terms across the three databases were grouped together under broader categories (Splicing, Extracellular matrix, Cell proliferation and tissue development, Ubiquitin related, Senescence, Cell Cycle, Receptor tyrosine kinases, Cell signalling, Cholesterol trafficking, ABC transporters, Calcium signalling, Aldosterone synthesis, Steroidogenesis, Cholesterol metabolism, and Other) based on shared genes.

### Visium (10x Genomics) spatial gene expression assay

Frozen adrenal gland samples (NAGs) were embedded in OCT (TissueTek) and cryo-sectioned (Thermo Scientific, CryoStar NX50) with a 10 µm thickness. The sections were then placed on tissue optimization and gene expression slides. 20 minutes was determined as the optimal permeabilisation temperature. Bright field H&E images were taken with a 10X objective (Leica DMi8 – A). After recovery of cDNA from the slide, the libraries were prepared following the gene expression user guide (10x Genomics – CG000239 Rev A). Subsequent to pooling, libraries were loaded at 200 pM and sequenced on a HiSeq-4000 (Illumina).

#### Generation of Visium count matrices

The base call (BCL) files generated from the raw sequencing data were converted to FASTQ reads using bcl2fastq tool (Illumina). The FASTQ reads were mapped to human reference dataset GRCh38 (build 2020-A; *refdata-gex-GRCh38-2020-A*) using Space Ranger v1.1.0 *count* pipeline. On average the two slides captured each 1269 spots under the tissue, and both yielded 1973 median genes and 225,346 mean reads per spot.

#### Seurat Processing of Visium count data

The two Visium Seurat objects were created using default parameters (*Load10X_Spatial*), and were subsequently normalised (*SCTransform*). Thirty PCs were computed and first 20 were used to generate the UMAP reduction. Knn of each spot was then determined (*FindNeighbors*) and clustering was performed (*resolution =0.2*, *FindClusters*), yielding 6 spatial clusters.

#### Label transfer from snRNA-seq data onto Visium

Label transferring was performed according to the label transfer workflow in Seurat. Anchors, or cell-spot pairwise comparisons were computed (*FindTransferAnchors*), and subsequently a prediction assay was generated (*TransferData*), containing the prediction scores for each cell type within each spot. Spatially variable features list were generated and included in both objects (*assay=“SCT”, selection.method=“markvariogram”*,*FindSpatiallyVariableFeatures*).

### Trajectory analysis

Both Visium objects were integrated with the SCTransform integration framework using default parameters. The Louvain clustering was performed (*resolution=0.2*) and those were annotated with prior knowledge from the label transfer analysis. The SCT adjusted count matrix (“counts” slot in SCT assay) was extracted from the integrated object, and a “cell data set” object was generated (*new_cell_data_set*) within Monocle 3^39^. Subsequently, the cds object was normalized (*preprocess_cds*) and seurat computed UMAP coordinates were added into reducedDims slot. The trajectory was then fit (*close_loop =F, use_partition =F*, *learn_graph*) and pseudotime was computed (*order_cells*) by setting the root to *RSPO3+* cells. After getting the genes that vary as a function of pseudotime (*neighbor_graph=“principal_graph”*, graph_test), they were grouped into modules (*resolution = 0.02, find_gene_modules*). Genes from top scoring modules (0 for Capsule, 1 for ZG and 2 for ZF_ZR) were combined to construct the top gene list that differentially change across pseudotime.

### ACA-NAG data integration

#### Count matrix merging and normalization

Count matrices from 12 ACA (37,037 nuclei: 7 CPA and 5 EIA) and highest scoring 5154 NAG nuclei (See Module Score section above) were merged. The Seurat object was then generated (*min.features=200, min.cells=3, CreateSeuratObject*), cell cycle (S.Score, G2M.Score) and mitochondrial genes were scored (*PercentageFeatureSet)* and regressed out during the normalisation step with SCTransform^91^.

#### Integration using Harmony

Post normalisation, sample integration was performed within the Harmony framework (1. *RunPCA, 2. RunHarmony with 40 PCs*)^92^. Using the Harmony dimensions, knns were computed (*FindNeighbors*) and clustering was done (*resolution=0.2*, *FindClusters*). Finally, a UMAP reduction was performed on the Harmony dimensions (*RunUMAP*). Clusters with less than 5 differentially expressed features were removed.

#### Analysis of DEG among the ACA clusters

Logistic regression test was used within the FindMarkers framework implemented in Seurat for computing DEGs across subtype (NAG, EIA, CPA) and clusters. The results were visualised as volcano plots (EnhancedVolcano v1.6.0) and heatmaps (ComplexHeatmap v2.4.3)

#### Cluster distribution among tissue samples

The distribution of the clusters among the samples were evaluated considering the type of the tissue (NAG vs EIA vs CPA) and the mutational status (no driver vs *CTNNB1*-mutation vs *PRKACA/GNAS*-mutation) for clusters selectively expressed only in ACA. To this aim, the percentage of the number of cells within the cluster *per* sample was considered. Data were analysed using the non-parametric Student’s T-tests and Mann-Whitney U test, as appropriate. A *p*-value <0.05 was considered statistically significant. The statistical analysis was performed with GraphPad Prism version 9 (GraphPad Software, San Diego, CA, USA).

### Immunohistochemistry

IHC was used to validate the expression and spatial context of the proteins of selected genes from the transcriptome analysis. This assay was performed with FFPE material from 16 NAGs, as previously described^93,94^. Primary antibodies are summarised in Supplementary Table 5. Moreover, double immunostaining was applied to validate new markers of adrenocortical stem/progenitors and adrenomedullary progenitors. The combination of NR2F2/ID1 and CHGA/SYT1 was used to test FFPE consecutive section from 9 NAGs. Details of the IHC assay including image acquisition is reported in the Supplementary material.

## Data availability

Raw gene expression counts can be found under “Supplementary Data” a the website accompanying this paper (https://……). Sequencing data was submitted to the ArrayExpress database of the European Bioinformatics Institute (EBI); accession number: E-MTAB-12129. Visium data can be retrieved from here: https://doi.org/10.5281/zenodo.5289292.

## Acknowledgements

The authors are grateful to Sonja Steinhauer, Antonia Lorey, and Martina Zink for excellent technical support. We are also thankful to Celso E. Gomez-Sanchez who provided selective antibodies detecting human CYP11B2. We thank Thomas Conrad for administrative and technical support.

This work has been supported by the Deutsche Forschungsgemeinschaft (DFG) (project FA-466/8-1, RO-5435/3-1 and 405560224 (M.F., C.L.R., and S.S.)) and within the CRC/Transregio (project number: 314061271 – TRR 205), and the Deutsche Krebshilfe (70113526 M.F. and C.L.R.). This work has been carried out with the help of the Interdisciplinary Bank of Biomaterials and Data of the University Hospital of Würzburg and the Julius Maximilian University of Würzburg (IBDW), supported by the Federal Ministry for Education and Research (Grant number FKZ: 01EY1102).

## Author contributions

C.L.R., M.F. and S.S. designed and coordinated the study and provided funding. B.A., L.S.L., and C.L.R. collected the tissues samples. A.K.S., P.A., S.N.V. and C.B. prepared the samples for the sequencing and generated sequencing data. A.K.S., So.Sa. and C.F. performed bioinformatics analysis. A.K.S. performed Visium experiments. B.A. and Si.Sb. performed immunohistochemistry experiments. S.H. evaluated the mutational status of the included adenomas. B.A. and A.K.S. performed statistical analysis and generated figures. Data interrogation and interpretation were carried out by B.A., A.K.S., So.Sa., C.F., Si.Sb., M.F., C.L.R., and S.S. B.A., A.K.S., C.L.R. and S.S. wrote the manuscript and Si.Sb., L.S.L. and M.F. provided critical feedback. All other authors read and approved the manuscript.

## Supplementary figures

**Supplementary Fig. 1.**
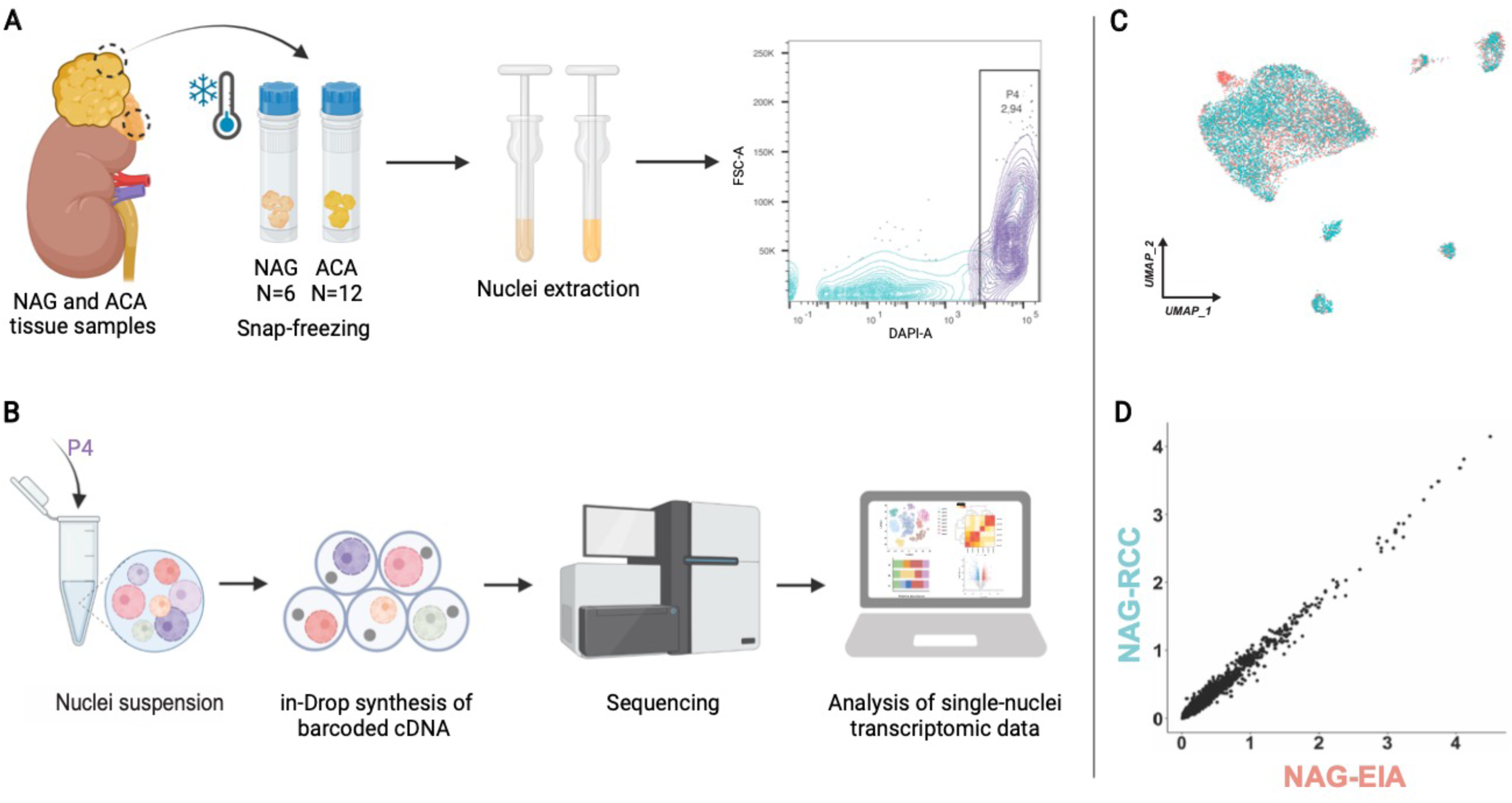

**Supplementary Fig. 2.**
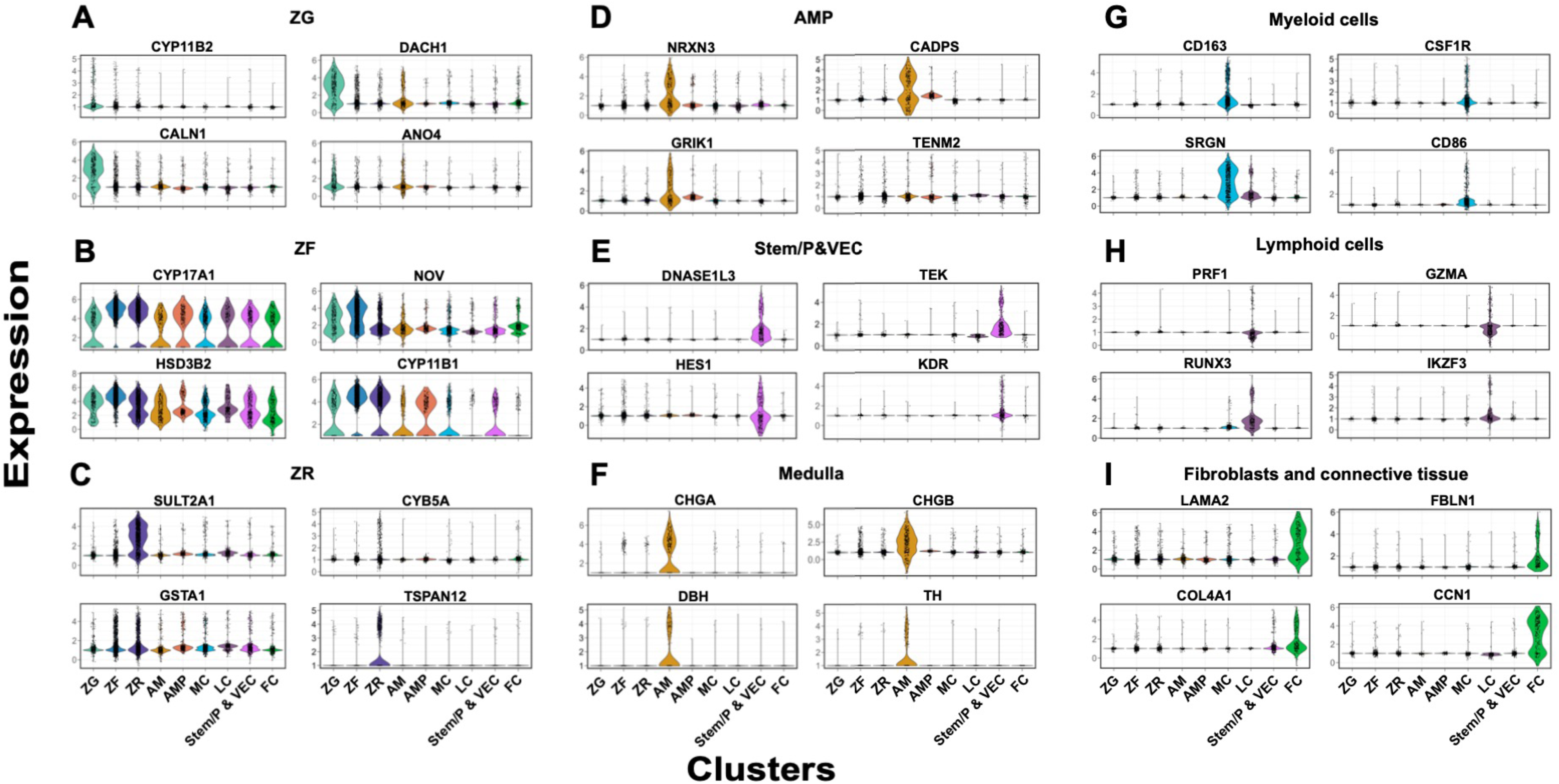

**Supplementary Fig. 3.**
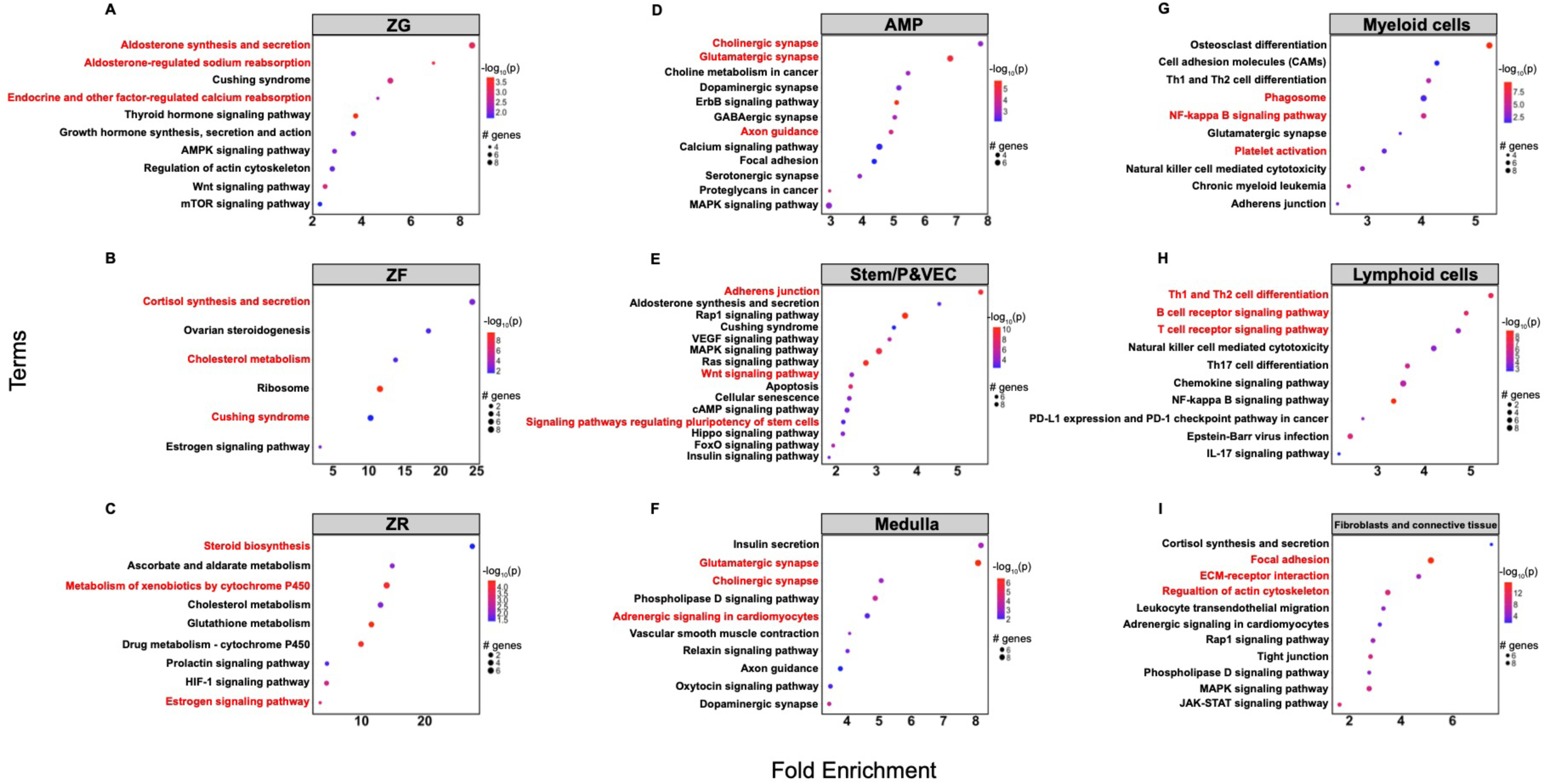

**Supplementary Fig. 4.**
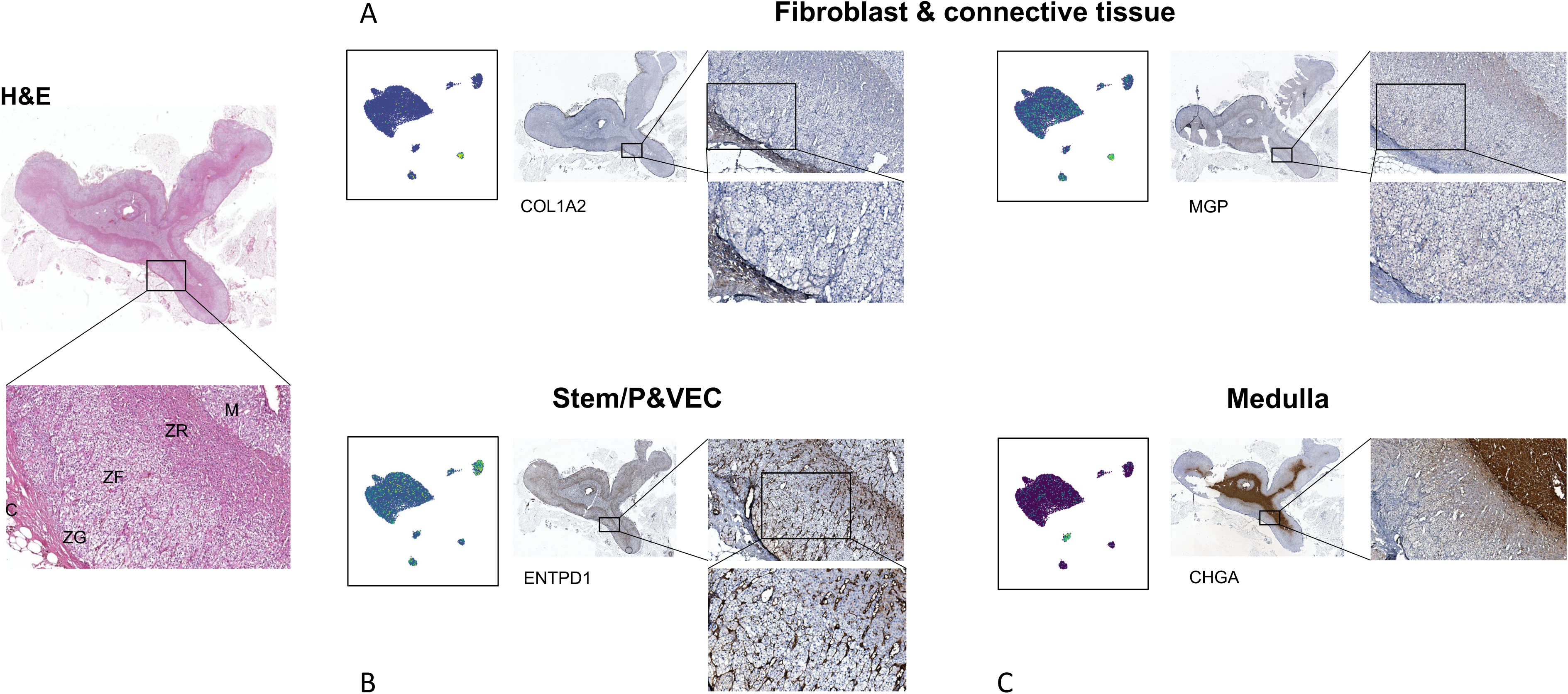

**Supplementary Fig. 5.**
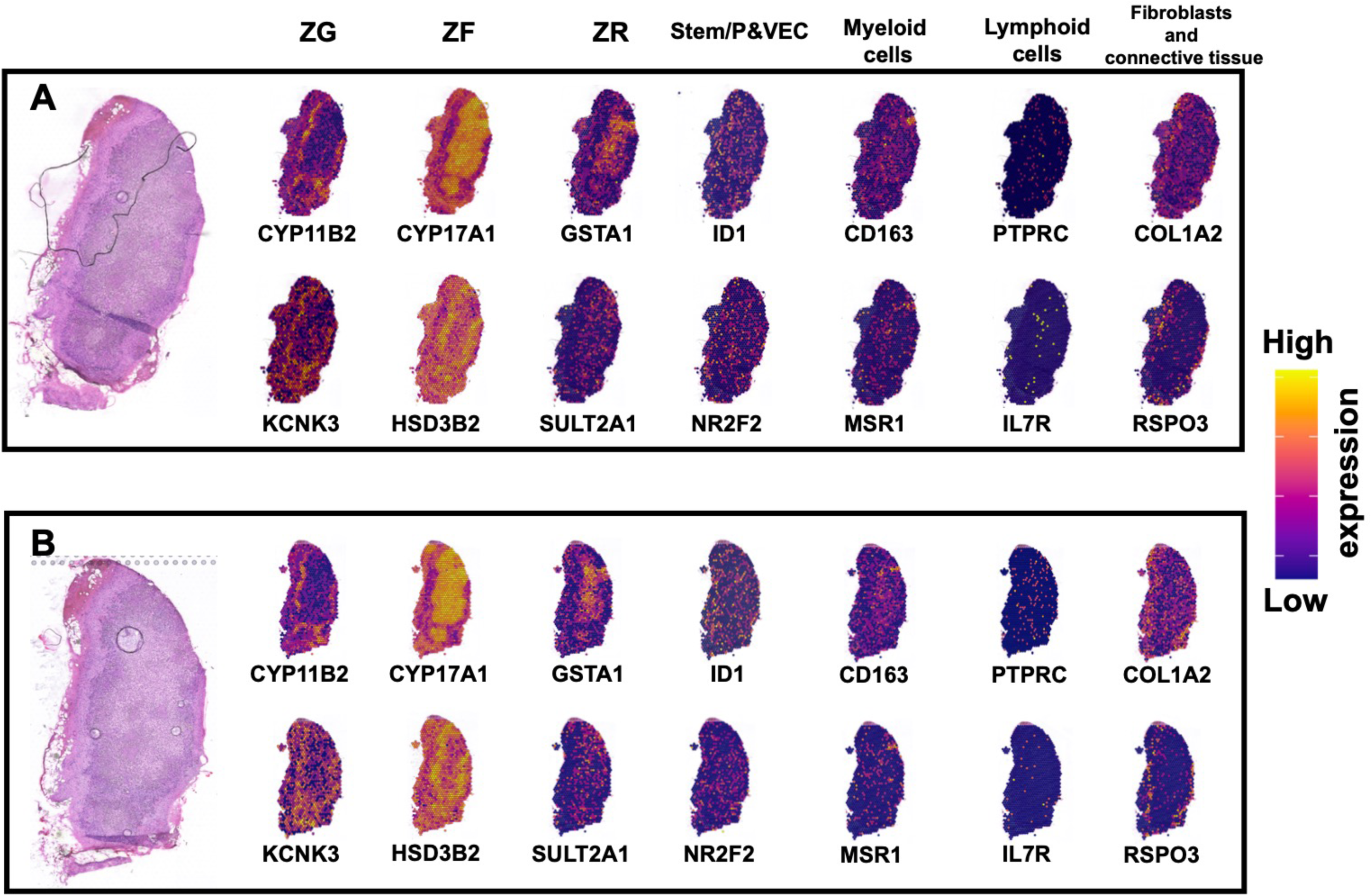

**Supplementary Fig. 6.**
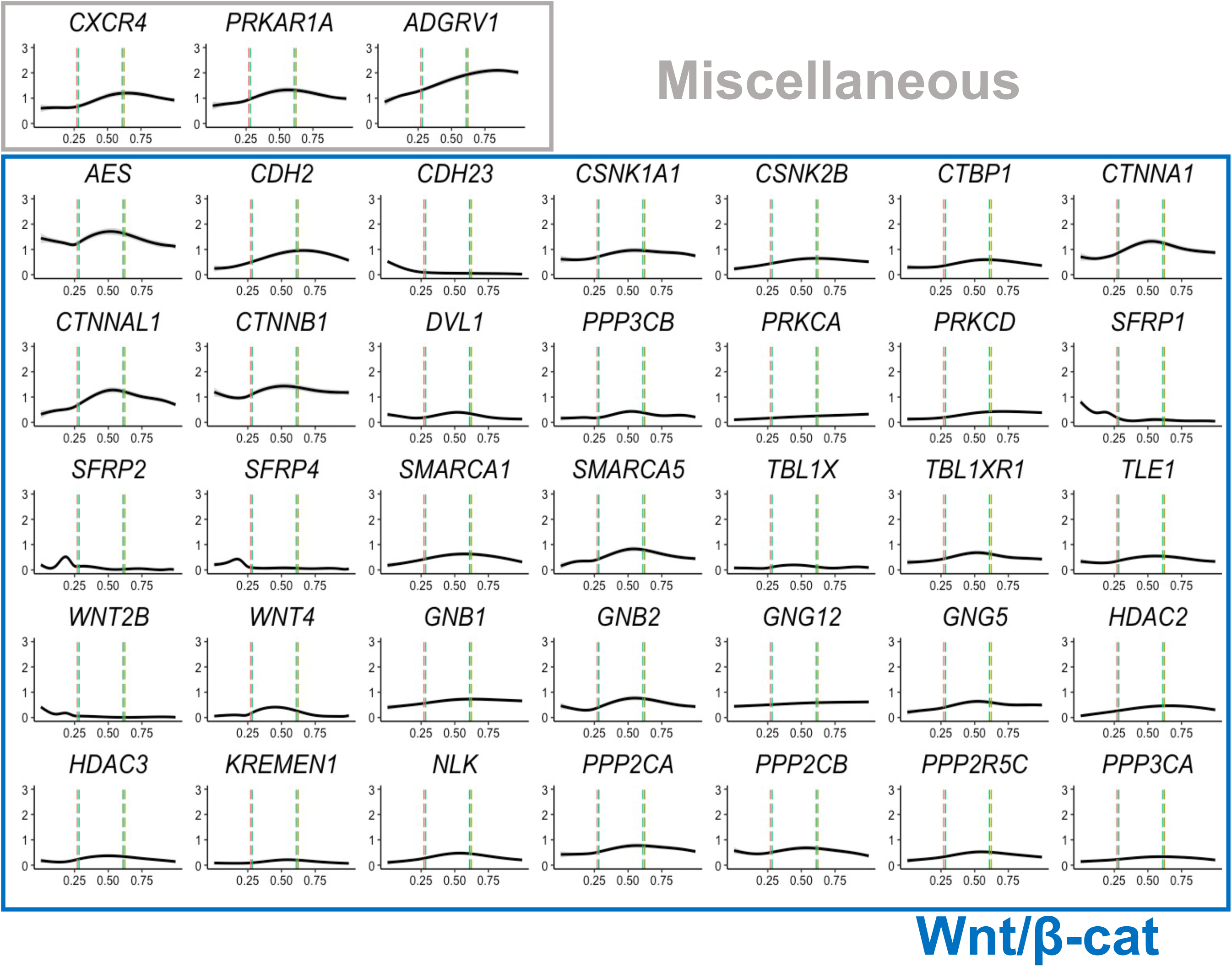

**Supplementary Fig. 7.**
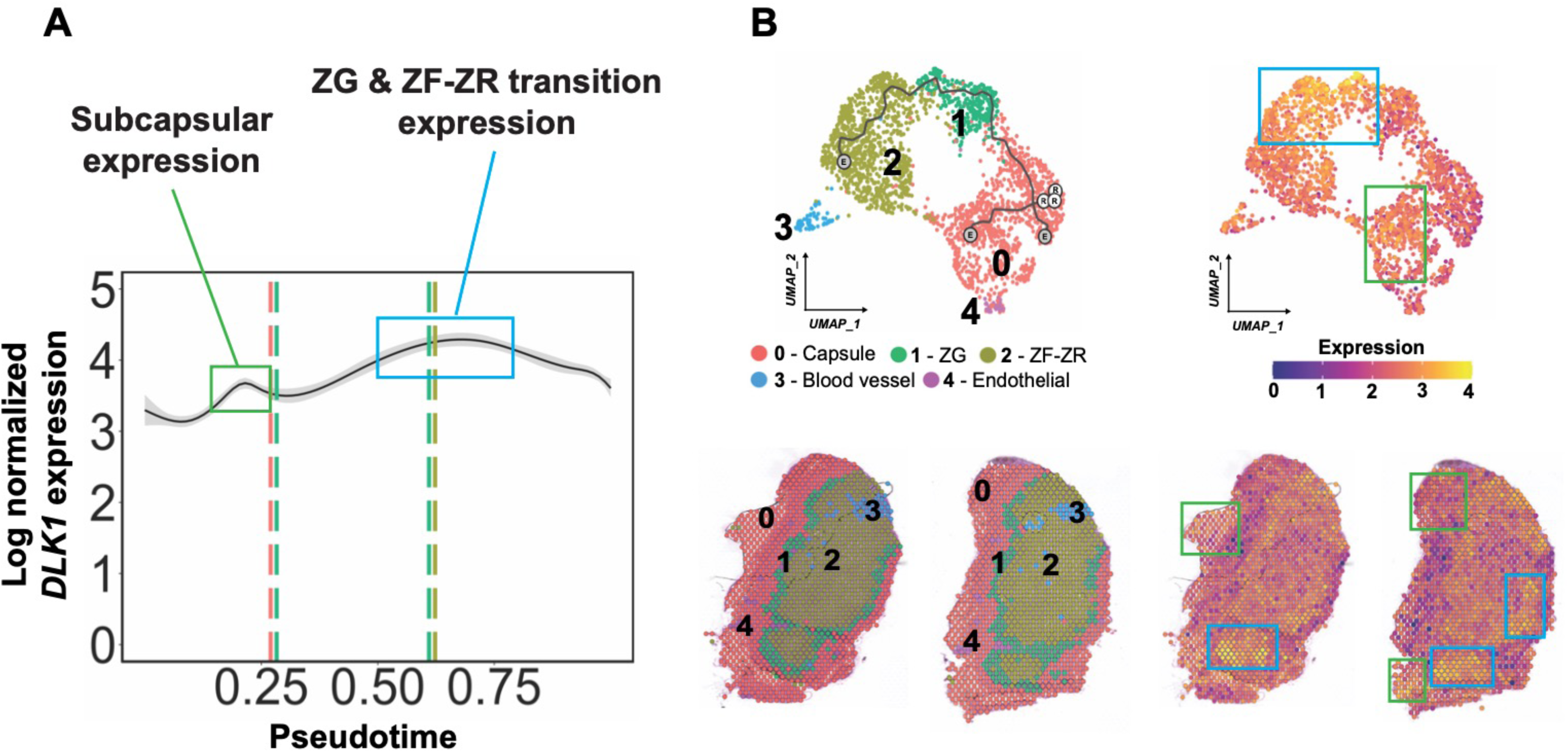

**Supplementary Fig. 8.**
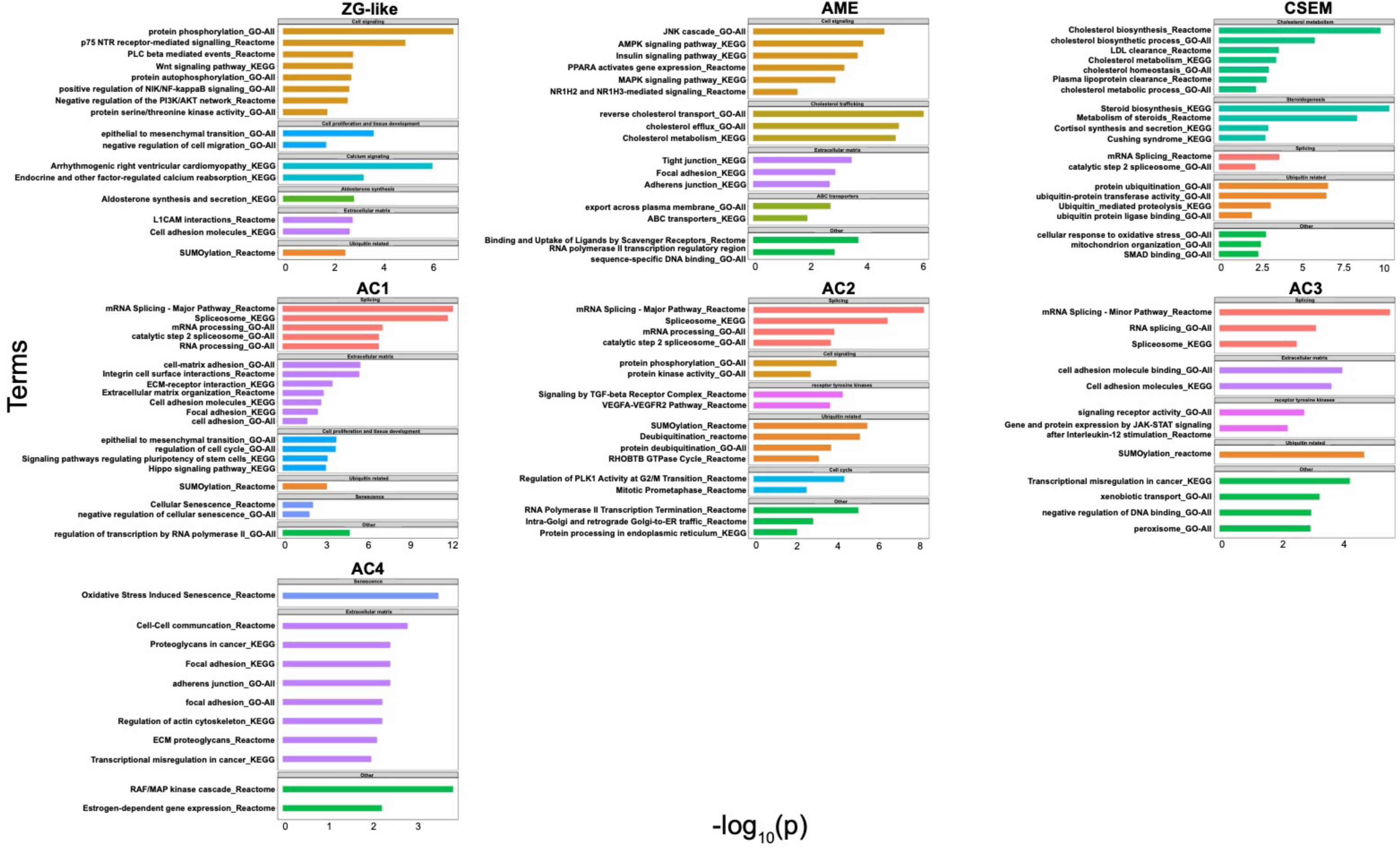

**Supplementary Fig. 9.**
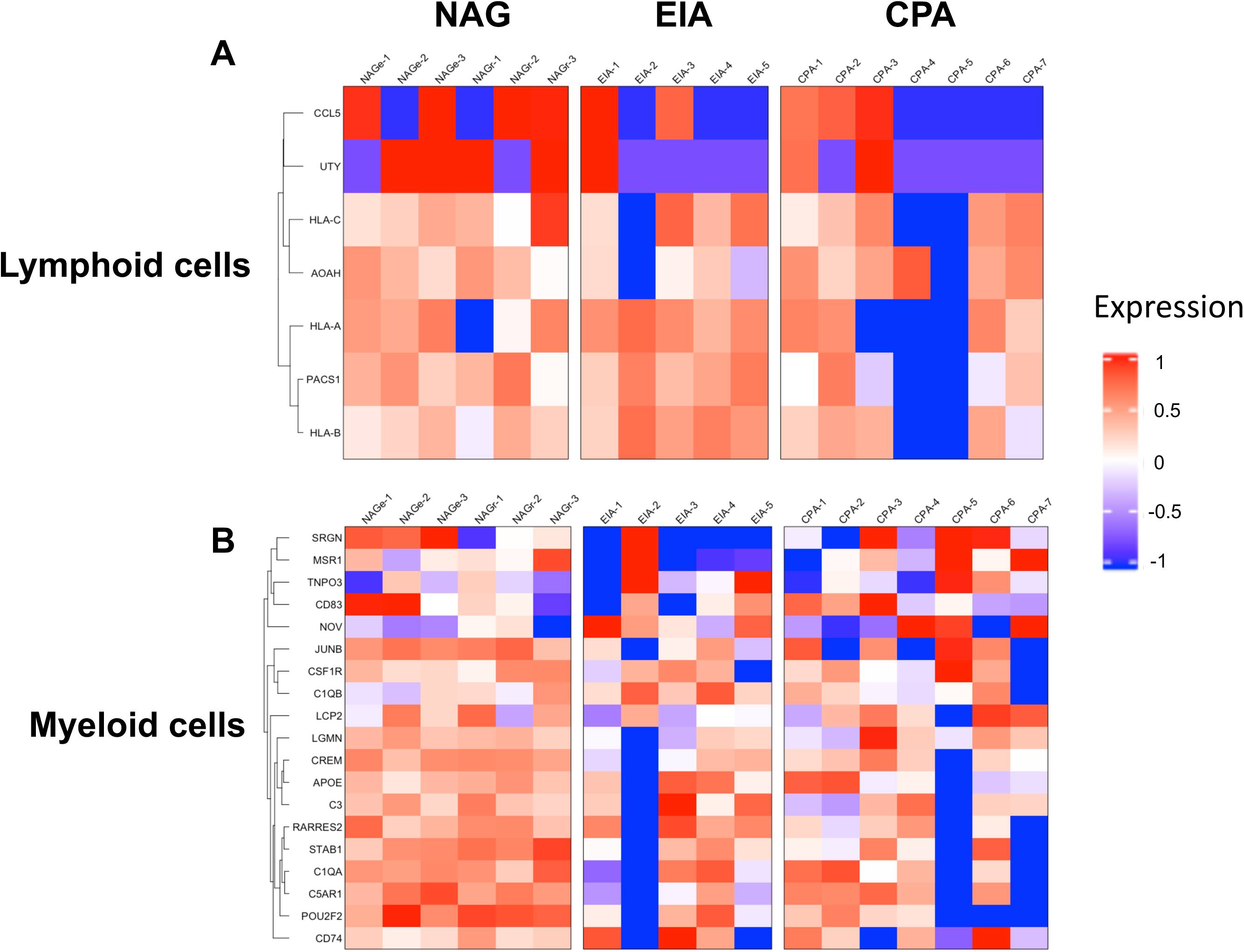

