## Supplemental Material for "Cell Atlas at Single-Nuclei Resolution of the Adult Human Adrenal Gland and Adrenocortical Adenomas"

**Supplementary Material**

**Supplementary Methods**

***Tissue sample collection:*** Subsequent to surgery, patient materials were collected and immediately stored at -80°C after flash freezing. The dissection of the adrenal tissues was made by expert pathologists[1]. The tissues with an average size of about 5x5x5 mm were ground gently using a mortar and a pestle in liquid nitrogen until obtaining a granular powder. Hereafter, half of the ground tissue was conserved at -80°C and the other half was used for downstream processing.

The aforementioned patient materials were selected from Würzburg Adrenal Biomaterial Archive, part of BMBF-funded Interdisciplinary Bank of Biomaterials and Data Würzburg (IBDW) applying the highest standards for biobanking[2].

***Clinical data collection:*** Hormone levels were measured using commercially available analytical procedures: serum cortisol and adrenocorticotropic hormone (ACTH) were analysed by Immulite 2000 Xpi from Siemens; late-night salivary cortisol (LNSC) was measured by a manual luminescence immunoassay from IBL; 24-h urinary free cortisol (UFC) was analysed by a manual radioimmunoassay from Immuntech, as previously reported[3].

The following screening tests for the diagnosis of cortisol-producing adenomas were applied as follows: 1 mg dexamethasone suppression test (DST) with a cutoff value of 50 nmol/l, LNSC (assay-specific reference normal range 0-4.1 nmol/L) and/or 24-h UFC (assay-specific reference normal range 22-193 nmol/d)[4].

***Single nuclei isolation:*** Ground snap-frozen adrenal tissue was homogenized in a Triton X-100 based lysis buffer (10% Triton X-100, Superasin 20 U $\mu L^{-1}$, RNaseIn U $\mu L^{-1}$, Nuclei Isolation Medium: 1 mM DTT, 50x Protease Inhibitor, 1.5M Sucrose, 1M KCl, 1M MgCl2, 1M Tris buffer pH 8.0). The homogenized nuclei were stained with DAPI (4',6-diamino-2-phenylindole) and sorted by FACS (BD FACSAria$^{TM}$ Fusion cell sorter, BD Genomics, USA) into RNAse inhibitor containing tubes (Supplementary Fig. 1).

***Immunohistochemistry***

Briefly, consecutive FFPE serial section of 16 NAGs were deparaffinized and rehydrated in descending graded series of ethanol. High-temperature antigen retrieval was achieved in 10 mM citric acid monohydrate buffer (pH 6.5) in a pressure cooker. Blocking of unspecific binding was performed with 20% human AB serum in PBS for 1 h at room temperature. Primary antibodies (summarized in Supplementary Table 5) were incubated 1 hour at room temperature. As negative control, tissue sections were incubated with N-Universal Negative Control Anti-Rabbit or Anti-Mouse (IS600 and IS750, respectively, Dako, Glostrup, Denmark), depending on the primary antibody used. Signal amplification was achieved by HiDef Detection HRP Polymer System (954D-50, Medac Diagnostika, Germany) followed by 10 min development with DAB substrate kit (957D-30, Cell Marque, USA). Nuclei counterstaining was obtained using with Mayer's haematoxylin for 2 min (T865.1, Carl Roth, Germany).

Double immunostaining was used to validate the new found markers of adrenocortical stem/progenitors and adrenomedullary progenitors. The combination of NR2F2/ID1 and CHGA/SYT1 was used. FFPE consecutive section form 9 NAGs were evaluated.

Blocking of unspecific binding was performed using 20% human AB serum in TBS solution. Slides were incubated with NR2F2 or CHGA antibodies and visualized in fuchsin-red by alkaline phosphatase MACH 3 Mouse AP-Polymer Detection (M3M532, Biocare Medical, CA, USA) and Vulcan Fast Red Chromogen Kit 2 (FR805H, Biocare Medical, CA, USA). A following incubation with secondary primary antibodies ID1 or SYT1 was performed. After signal amplification and detection with HiDef Detection HRP Polymer System, ID1 or SYT1 were stained in blue-green by Vina Green Chromogen (BRR807AH, Biocare Medical, CA, USA). Nuclei counterstaining was obtained using with Mayer's haematoxylin for 1 min.

All images were acquired by Leica Aperio Versa Brightfield scanning microscope (Leica, Germany) using same light intensity parameters to avoid biased information. A 1x, 10x and 20x enlargement were used to evaluate the staining.

**Supplementary Table 1. Top 100 differentially up-regulated genes among the clusters of adult human normal adrenal glands.**

List of top hundred differentially expressed genes (DEGs) among the different clusters of the normal adrenal glands. Clustering was done at *resolution=0.01* using the *FindClusters* function (as detailed in Methods).

**Supplementary Table 2. Top 100 differentially up-regulated genes among the clusters of adult human adrenocortical adenomas.**

List of top hundred differentially expressed genes (DEGs) among the different clusters of adrenocortical adenomas. Clustering was done at *resolution=0.01* using the *FindClusters* function (as detailed in Methods).

**Supplementary Table 3. Details on clinical findings, imaging and histological characteristics of the adrenocortical adenomas.**

Abbreviation: AC1-4, adenoma cluster 1-4; CSEM, cholesterol-and steroid-enriched metabolism; CT, computed tomography; EIA, endocrine inactive adenoma; FDG-PET/CT, fluorodeoxyglucose (FDG)-positron emission tomography (PET)/computed tomography (CT); HPF, high-power field; HU, Hounsfield unit; na, not available; RNA-Seq, RNA-sequencing (data from previous study[1]); WES, whole exome sequencing (data from previous study[5])

**Supplementary Table 4. Summary of normal adult adrenal glands (NAG) evaluated by immunohistochemistry.**

Abbreviation: EIA, endocrine inactive adenoma; F, female; M, male; n.a., not available; NAGe, normal adrenal gland from the tissue surrounding EIA; NAGr, normal adrenal gland from adrenalectomies performed during surgery for RCC; RCC, renal cell carcinoma

**Supplementary Table 5. Primary antibodies used for the immunohistochemistry.**

**Supplementary Table 6. PCR primer sequences used for detecting the hot-spot mutation in *CTNNB1, PRKACA and GNAS*.**

Known drivers hot-spot mutations in *CTNNB1* (entire exon 3), *PRKACA* (p.Leu206 in exon 7) and *GNAS* (p.Arg201 and p.Gln227) [1,6] were evaluated by Sanger sequencing. Abbreviation: PCR, Polymerase Chain Reaction.

**Supplementary Fig 1. Sample preparation workflow.**

**A.** Six normal human normal adrenal glands (3 NAGs from endocrine inactive adenoma "NAG-EIA" and 3 from renal cell carcinoma "NAG-RCC" patients) and 12 adrenocortical adenoma (ACAs: 5 from EIA and 7 from cortisol producing adenoma (CPA) patients) samples were collected and snap-frozen. The dissection of the adrenal gland was done by an expert pathologist. Nuclei extraction was performed via dounce homogenization[7]. Using FACS, nuclei were sorted into tubes via DAPI discrimination. **B.** Subsequently, the samples were processed with the inDrop**™** (1CellBio) system, according to the manufacturer's protocol. **C.** UMAP representation of the single-nuclei transcriptomes from the six normal samples (NAG-EIA in red; NAG-RCC in blue). **D.** Average gene expression correlation between NAG-EIA and NAG-RCC patients (Pearson correlation coefficient (r = 0.99).

**Supplementary Fig 2. Violin plots representing cluster-specific gene expression in adult human normal adrenal glands at single-nuclei transcriptome level.**

**A-C:** Subclusters of the normal adrenal cortex: ZG, ZF and ZR. **D-I:** satellite clusters of the normal adrenal cortex, namely the AMP, Stem/P&VEC, Medulla, Myeloid cells, Lymphoid cells, Fibroblasts and connective tissue.

**Supplementary Fig 3. Gene set enrichment analysis of single-nuclei transcriptome in adult human normal adrenal glands.**

Gene set enrichment analysis was performed using pathfindR (KEGG) and results for each cluster is represented separately. The dot size varies with the quantity of observed significant genes that constitute their representative enriched term. The colour scale represents the –log10(p) value. The x-axis indicates fold enrichment, while the y-axis covers the enriched terms within the representative clusters.

**Supplementary Fig 4. Validation of most relevant markers of the Fibroblasts & connective tissue, Stem/P&VEC and Medulla clusters by immunohistochemistry (IHC).**

Hematoxylin- and Eosin- (H&E) staining served to validate the different zones of the normal adrenal gland, precisely all three zones of the cortex (Zona glomerulosa, ZG; fasciculata, ZF; reticularis, ZR), together with the capsule (C) and the adrenal medulla (M) were observed. For the IHC assay, **A.** COL1A2 and MGP were used as protein markers of the fibroblasts & connective tissue cluster (FC); **B.** ENTPD1 as protein marker for vascular endothelial cells (Stem/P&VEC); and **C.** CHGA as protein marker for the medulla (AM). All images were acquired by Leica Aperio Versa brightfield scanning microscope (Leica, Germany). A 1x picture with an enlargement of 10x of the same area of consecutive slides was used to generate IHC photos. In some cases, an enlargement of 20x was used to better show the different distribution of the stained proteins in the adrenal glands.

**Supplementary Fig 5. Spatial gene expression plots.**

Spatial gene expression plots in both replicates, representing expressed genes specific for different clusters: ZG, ZF, ZR, Stem/P&VEC, Myeloid cells, Lymphoid cells, Fibroblasts and connective tissue.

**Supplementary Fig 6. Expression of WNT/ß-cat pathway genes across pseudotime.** Smoothed lines are generated based on the scatter profile of each gene (95% confidence interval displayed in grey around the line). Vertical lines represent the transition zones from 0-Capsule (red) to 1-ZG (green) and 1-ZG to 2-ZF-ZR (ochre). WNT/ß-cat genes are highlighted within the blue box while miscellaneous genes in grey.

**Supplementary Fig 7. *DLK1* expression across pseudotime**

**A.** Scatter plot representing variation of log normalized expression of DLK1 throughout pseudotime (its expression is highlighted by a green box in the subcapsular region and by a blue box at the ZG to ZF-ZR transition zone). Vertical lines represent the transition zones from 0-Capsule (red) to 1-ZG (green) and 1-ZG to 2-ZF-ZR (ochre). **B**. First row: Integrated UMAPs (Uniform Manifold Approximation and Projection) from both Visium sections (10X Genomics) showing the pseudotime trajectory (indicated by black line) and DLK1 expression. Second row: Visium sections showing the previously acquired label transferring clustering as well as DLK1 expression.

**Supplementary Fig 8. Gene set enrichment analysis of 7 newly identified clusters in ACA**

Gene set enrichment analysis was performed using pathfindR, using three databases (KEGG, Reactome, GO-All). Curated lists of terms are represented for each of the 7 clusters that were identified in ACA: similar terms were placed (Splicing, Extracellular matrix, Cell proliferation and tissue development, Ubiqutin related, Senescence, Cell signalling, receptor tyrosine kinases, Cell Cycle, Cholesterol trafficking, ABC transporters, Cholesterol metabolism, Steroidogenesis, Calcium signalling, Aldosterone synthesis, Other) in groups based on their shared gene lists.

**Supplementary Fig 9. Comparison of the immune transcriptome across subtypes**

Heatmap representing DEGs across subtype and samples **A.** Lymphoid cells and **B.** Myeloid cells.

Hierarchical clustering was performed on the genes (rows) and the average expression values were scaled accordingly.
